# Rapid DNA Replication Origin Licensing Protects Stem Cell Pluripotency

**DOI:** 10.1101/164368

**Authors:** Jacob Peter Matson, Raluca Dumitru, Phillip Coryell, Ryan M Baxley, Weili Chen, Kirk Twaroski, Beau R. Webber, Jakub Tolar, Anja-Katrin Bielinsky, Jeremy Purvis, Jeanette Gowen Cook

**Author notes:** Corresponding Author: J.G.C.

## Abstract

Complete and robust human genome duplication requires loading MCM helicase complexes at many DNA replication origins, an essential process termed origin licensing. Licensing is restricted to G1 phase of the cell cycle, but G1 length varies widely among cell types. Using quantitative single cell analyses we found that pluripotent stem cells with naturally short G1 phases load MCM much faster than their isogenic differentiated counterparts with long G1 phases. During the earliest stages of differentiation towards all lineages, MCM loading slows concurrently with G1 lengthening, revealing developmental control of MCM loading. In contrast, ectopic Cyclin E overproduction uncouples short G1 from fast MCM loading. Rapid licensing in stem cells is caused by accumulation of the MCM loading protein, Cdt1. Prematurely slowing MCM loading in pluripotent cells not only lengthens G1 but also accelerates differentiation. Thus, rapid origin licensing is an intrinsic characteristic of stem cells that contributes to pluripotency maintenance.

## INTRODUCTION

Metazoan DNA replication requires initiation at thousands of DNA replication origins during S phase of every cell cycle. Origins are genomic loci where DNA helicases first unwind DNA and DNA synthesis begins. Origins are made competent for replication during G1 phase of each cell cycle by the loading of Minichromosome Maintenance (MCM) complexes onto DNA. MCM is the core component of the replicative helicase, and the process of MCM loading is termed origin licensing. Total MCM levels remain constant throughout the cell cycle, but the levels of DNA-loaded MCM change as cells progress through the cell cycle. Cells can begin MCM loading as early as telophase, and loading continues throughout G1 until the G1/S transition, the point of maximum DNA-loaded MCM^1,2^. Throughout S phase, individual MCM complexes are activated for DNA unwinding as origins “fire”. MCM complexes travel with replication forks and are progressively unloaded as replication forks terminate (Fig. 1a)^3–5^.

**Figure 1.**
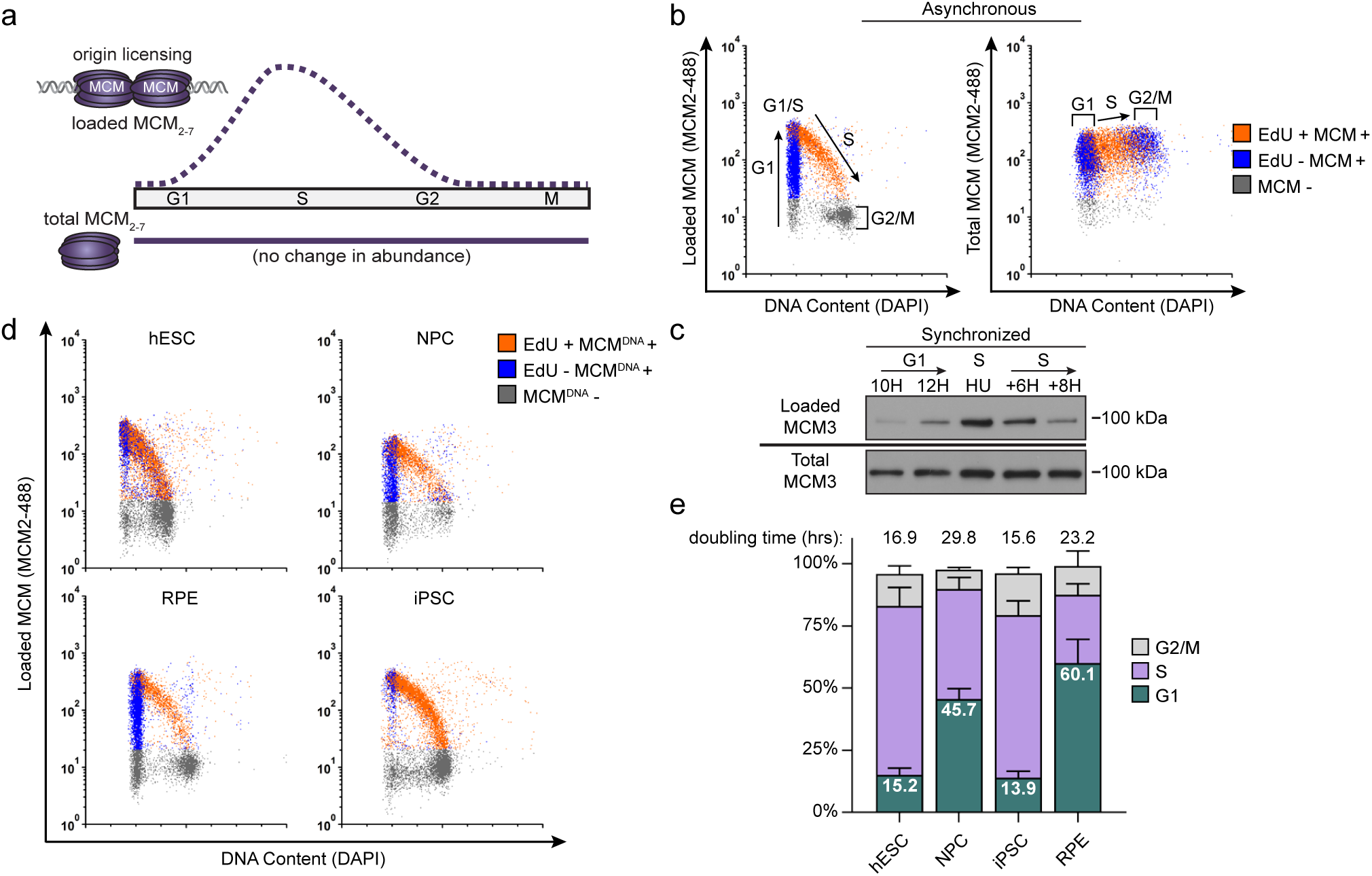
Pluripotent stem cells load MCMs faster than differentiated cells. (a) DNA-loaded MCM levels increase in G1 and decrease in S phase whereas total MCM protein levels are constant throughout the cell cycle. (b) Flow cytometric analysis of DNA-loaded and total MCM in asynchronously proliferating RPE1-hTERT cells. Cell cycle phases are defined by DNA content and DNA synthesis. Left: Cells were labeled with EdU, extracted with nonionic detergent to remove unbound MCM, fixed, and stained with anti-MCM2 (a marker for the MCM_2-7_ complex), DAPI (total DNA), and for EdU incorporation (active DNA synthesis). Right: Cells were treated as on the left except that they were fixed prior to extraction to detect total MCM2. (c) T98G cells were synchronized in G0 by contact inhibition and serum deprivation, then released into G1 for 10 or 12 hrs, or re-synchronized in early S with hydroxyurea (HU), and released into S for 6 or 8 hrs. MCM3 in chromatin-enriched fractions (Loaded) or whole cell lysates (Total) was detected by immunoblotting. (d) Chromatin flow cytometry of the indicated asynchronous cell lines measuring DNA content (DAPI), DNA synthesis (EdU incorporation), and loaded MCM (anti-MCM2). Blue Cells are MCM^DNA^-positive and EdU-negative (i.e. G1), orange are both MCM^DNA^-positive and EdU-positive (S phase); grey are MCM^DNA^-negative. (e) Stacked bar graph of cell cycle phase distribution from cells in (d); mean with error bars ± SD (n=3 biological replicates). The doubling times were calculated experimentally using regression analysis in GraphPad Prism.

The control of origin licensing is critical for genome stability. Origins must not be re-licensed after S phase begins because such re-licensing can cause a genotoxic phenomenon known as re-replication which may result in double strand breaks, gene amplification, aneuploidy, and general genome instability^6,7^. To avoid re-replication, MCM loading is tightly restricted to G1 phase by multiple overlapping mechanisms that destroy or inactivate MCM loading proteins to prevent any new origin licensing after S phase begins^4,6,7^. On the other hand, cells typically load 5-10-fold more MCM complexes in G1 than they strictly require under ideal circumstances, and the additional MCM loading ensures timely and complete genome duplication even if replication hurdles are encountered in S phase^8–10^. It is possible for cells to proliferate with less than optimal MCM loading, but such cells are hypersensitive to DNA damage and replication stress^11,12^.

MCM loading to license origins is restricted to G1, but G1 length varies widely among different cell types. For example, specialized developmental and immune cell cycles have minimal G1 lengths of mere minutes^13–15^. In cultured human embryonic stem cells, G1 is only 2-3 hours, and this short G1 is both a hallmark of and has been implicated in maintaining pluripotency^16,17^. G1 lengthens early in differentiation, and in cultured somatic cells is often greater than 12 hours^18^. Thus, different proliferating cells have drastically different amounts of time available to complete MCM loading before making the G1-to-S phase transition. In addition, pluripotent stem cells respond to differentiation stimuli specifically in G1 phase, suggesting that the balance among cell cycle phases influences differentiation potential^19,20^.

Given that MCM loading is restricted to G1 and the wide variation of G1 lengths, we postulated that the absolute amount of loaded MCM in S phase is a product of both the time spent in G1 and the rate of MCM loading. The combination of these two parameters has implications for genome stability because loading more or less MCM in G1 influences S phase length and how effectively S phase cells can accommodate both endogenous and exogenous sources of replication stress^21,22^. These implications are relevant both when cell cycle distributions change during development and during oncogenesis since many cancer cell lines also have short G1 phases. The actual rate of MCM loading in human cells has not yet been quantified however, and little is known about how the amount, rate or timing of MCM loading varies between cells with different G1 lengths. Here we utilized single cell flow cytometry to measure MCM loading rates in asynchronous populations of pluripotent and differentiated cells. We discovered that rapid MCM loading is intrinsic to pluripotency, slows universally during differentiation, and rapid replication licensing suppresses differentiation. These findings demonstrate that the rate of MCM loading is subject to developmental regulation, and we suggest that rapid origin licensing is a new hallmark of pluripotency.

## RESULTS

### Pluripotent cells load MCM significantly faster than differentiated cells

We considered two possibilities for how cells with varying G1 lengths load MCM onto DNA. One possibility is that cells with short G1 phases load MCM at the same rate as cells with long G1 phases resulting in less total loaded MCM. Alternatively, cells with short G1 phases could load MCM faster than cells with long G1 phases and reach similar levels of loaded MCM. To distinguish between these scenarios, we developed an assay to measure DNA-loaded MCM in individual cells of asynchronously proliferating populations by adapting a previously-reported flow cytometry method^23,24^. We extracted EdU-labelled immortalized epithelial cells (RPE1-hTERT) with nonionic detergent to remove soluble MCM. We then fixed the remaining chromatin-bound proteins for immunofluorescence with anti-MCM2 antibody as a marker of the MCM_2-7_ complex (Fig. 1b, left) or without primary antibody as a control (relevant flow cytometry gating schemes are shown in Supplementary Fig. 1). Interestingly, individual G1 cells have a very broad range of DNA-loaded MCM levels with a more than 100-fold difference between minimum and maximum (Fig. 1b, left). MCMs are unloaded during S phase, ending in G2/M with undetectable MCM on DNA (Fig. 1b, left). As expected, total MCM protein levels do not substantially change in the cell cycle (Fig. 1b, right). For comparison to commonly used cell fractionation methods to assess MCM dynamics, we also probed immunoblots of chromatin-enriched fractions, and noted a similar MCM expression, G1 loading, and S phase unloading pattern (Fig. 1c)^25,26^.

Loaded MCM complexes are extremely stable on DNA, both *in vivo* and *in vitro*^27–30^. In human cells, MCMs can persist on DNA for more than 24 hours during a cell cycle arrest and are typically only unloaded during S phase^31^. These properties result in MCM loading that occurs unidirectionally throughout G1 phase^32^. The unidirectional nature of MCM loading means that G1 cells with low MCM levels are in early G1, and G1 cells with high MCM levels are in late G1. Since we observed a broad distribution of MCM loading throughout G1 including many cells with low levels of loaded MCM, we conclude that RPE1-hTERT cells load MCM relatively slowly during their ∼9 hr G1.

We then used this method to analyze MCM loading in asynchronous cells with different G1 lengths. H9 human embryonic stem cells (hESCs) have a short G1 phase and spend most of the cell cycle in S phase. In contrast to the differentiated epithelial cells, the majority (∼80%) of G1 hESCs had high levels of loaded MCM; very few G1 cells had low levels of loaded MCM (blue cells, Fig. 1d). This difference suggests that hESCs load MCM rapidly to achieve abundant DNA-loaded MCM in a short time. To test if MCM loading varies in differentiated cells, we differentiated hESCs into neural progenitor cells (NPCs) to generate an isogenic pair of pluripotent and differentiated cells. In contrast to hESCs, differentiated NPCs had a wide distribution of DNA-loaded MCM in G1 (blue cells, Fig. 1d); they also spend approximately five time longer in G1 (Fig. 1e). Since the NPCs had many cells with low levels of DNA-loaded MCM, we conclude that these differentiated cells load MCM more slowly than hESCs.

To generate another isogenic pair of pluripotent and differentiated cells we reprogramed ARPE-19 primary retinal pigmented epithelial cells (RPE) into induced pluripotent stem cells (iPSCs). The iPSCs had hallmark features of pluripotency as measured by microscopy, bisulfite sequencing, gene expression, and teratoma formation (Supplementary Fig. 2a-e), and their G1 phases were typically seven times shorter than their differentiated parents (Fig. 1e). Like hESCs, the pluripotent iPSCs had predominantly high levels of DNA-loaded MCM in G1 (Fig. 1d). Importantly, both of the pluripotent cell lines reached approximately equal levels of DNA-loaded MCM at the start of S phase as their differentiated counterparts did, but in in less time (the absolute MCM loading intensities are comparable when samples are processed and analyzed with identical instrument settings). Taken together, these data demonstrate that pluripotent cells load MCMs rapidly in G1, but differentiated cells load MCMs slowly.

We then quantified the relative MCM loading rates in pluripotent and differentiated cells using ergodic rate analysis, a mathematical method that can derive rates from fixed, steady state populations^33^. Ergodic analysis can measure any unidirectional rate parameter from a steady state distribution, and is not limited to the cell cycle (e.g. car traffic jams)^34^. The ergodic analysis as applied to the cell cycle means that within a steady state population with a constant doubling time and cell cycle distribution, the number of cells at any point in the cell cycle is inversely related to the rate they move through that point. For any measured parameter, the density of cells indicates rate: low cell density on a flow cytometry plot indicates a fast rate passing through that cell cycle state whereas high cell density indicates a slow rate. This phenomenon is analogous to a high density of slow-moving cars observed at a given point on a road in a traffic jam compared to a low density of fast-moving cars on an open highway. We visualized MCM loading as histograms of the MCM^DNA^ intensities in only the G1 cells for ergodic rate analysis (G1-MCM^DNA^, Fig. 2a, b and Supplementary Fig. 3).

**Figure 2.**
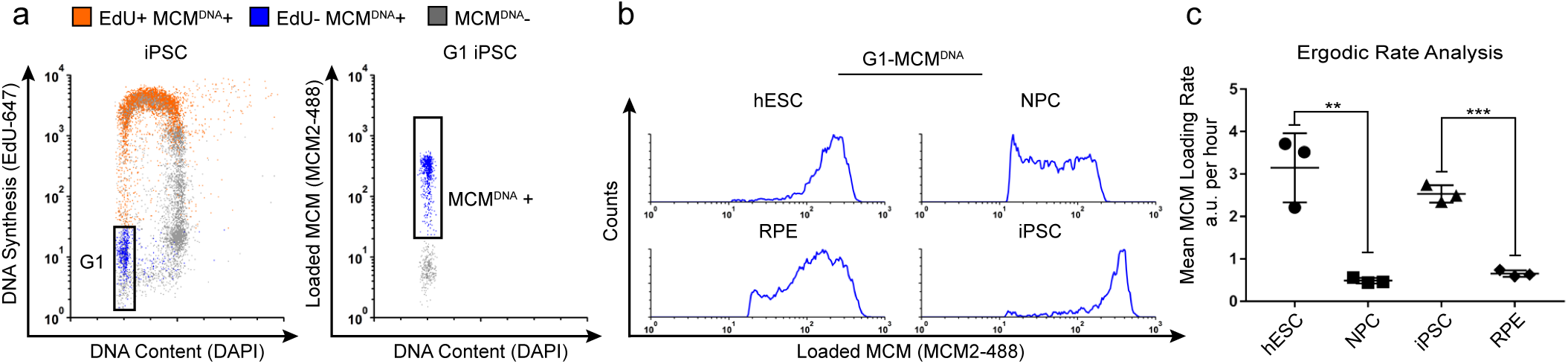
Quantification of MCM loading rate by ergodic rate analysis. (a) Gating scheme for chromatin flow cytometry of iPSCs measuring DNA content (DAPI), DNA synthesis (EdU incorporation), and loaded MCM (anti-MCM2); this sample is from Fig 1d. (b) Histograms of only the G1-MCM^DNA^ –positive cells from the four chromatin flow cytometry samples in Fig 1d. (c) Calculated mean MCM loading rate by ergodic rate analysis; mean with error bars ± SD. (n=3 biological replicates), unpaired two tailed t-test. **p=0.0049. ***p=0.001. See Methods for details.

To compute MCM loading rate per hour, we experimentally determined the cell cycle distributions and doubling times of each cell population (Supplementary Fig. 3). Pluripotent cells reached near equal levels of loaded MCM at the G1/S transition in less time than differentiated cells. To quantify the actual MCM loading rate difference, we subdivided the G1-MCM^DNA^ population into 10 equally-sized bins, calculated the MCM loading rate for each bin, then the overall average MCM loading rate for each G1 population. These calculations revealed that pluripotent hESCs loaded MCM 6.5 times faster than differentiated NPCs and pluripotent iPSCs loaded MCM 3.9 times faster than differentiated RPEs (Fig. 2c). Thus, pluripotent cells with short G1 phases load MCMs significantly faster than differentiated cells with long G1 phases.

### Differentiation, G1 length, and MCM loading rate are coupled

We hypothesized that MCM loading is fundamentally linked to pluripotency because MCM loading rate decreased during differentiation and increased during reprogramming. This idea predicts that slowed MCM loading is a phenomenon common to differentiation towards all germ layers. To test that hypothesis, we initiated differentiation in hESCs towards the three embryonic germ layers (neuroectoderm, mesoderm and endoderm) and extraembryonic trophectoderm, collecting cells at 24 hour and 48 hours after inducing differentiation (Supplementary Fig. 4b). We confirmed progress towards each lineage by the expected gene expression changes, particularly induction of lineage-specific markers and modest reduction of pluripotency markers – even at these very early time points (Fig. 3c). We assessed MCM loading rates during differentiation by flow cytometry as before. The MCM loading rate clearly decreased for all germ layers rapidly within the first 48 hours of initiating differentiation (Fig. 3a, compare the grey histograms for undifferentiated G1 cells to the green and blue histograms). The decrease in MCM loading rate also coincided with the increase in the proportion of G1 cells for each lineage. For example, within 24 hours of neuroectoderm differentiation, G1 had already lengthened and MCM loading had slowed, but during trophectoderm differentiation both G1 lengthening and slowed MCM loading took 48 hrs (Fig. 3a and b). The closely-coordinated changes that we universally observed during differentiation suggest that MCM loading rate is coupled to G1 length. Importantly, these results demonstrate that the rate of origin licensing by MCM loading is developmentally regulated.

**Figure 3.**
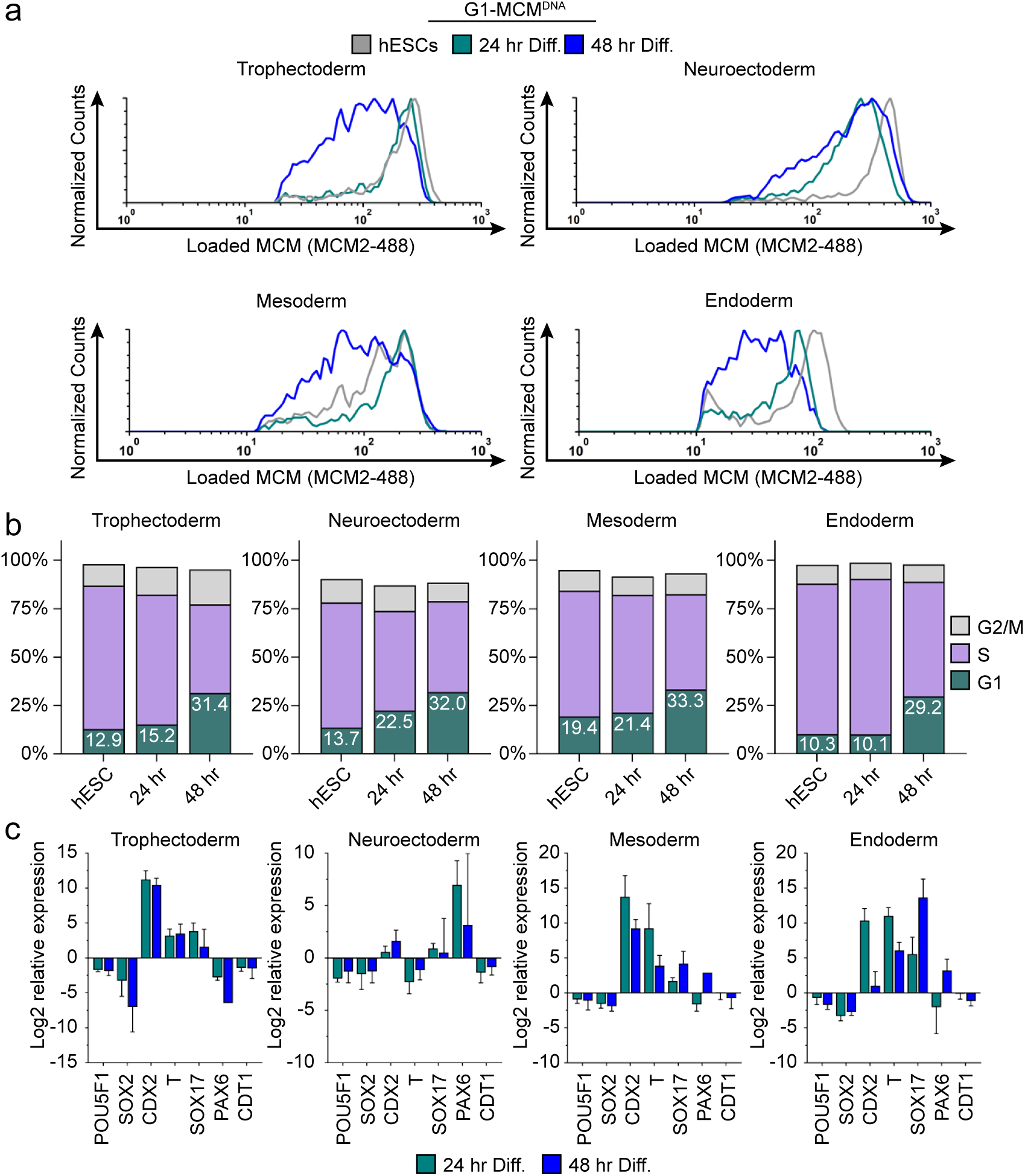
Differentiation universally decreases MCM loading rate. (a) Chromatin flow cytometry of hESCs induced to differentiate towards trophectoderm, neuroectoderm, mesoderm, or endoderm for 24 or 48 hrs. Histograms show only G1-MCM^DNA^ cells as in Figure 2b. See methods for differentiation protocols. Cell counts for 24 hr and 48 hr were normalized relative to corresponding hESC samples. (b) Stacked bar graphs of cell cycle distribution for cells in (a). (c) Gene expression analysis of differentiation markers by quantitative PCR of the samples in 3a; log^2^ expression is relative to the undifferentiated cells. Data are mean ±SD of two biological replicates.

We next asked if G1 length and MCM loading rate are *obligatorily* coupled, or if the link can be short-circuited by artificially advancing the G1/S transition. To distinguish between these possibilities, we constructed an RPE1-hTERT derivative with a *CYCLIN E1* cDNA under doxycycline-inducible control. Cyclin E1 overproduction reproducibly shortened G1 length, consistent with previous studies (Fig. 4a, b)^35,36^. Strikingly, cells overproducing Cyclin E1 (designated as “↑Cyclin E1”) not only spent less time in G1 but also began S phase with much lower amounts of loaded MCM compared to the control; this new subpopulation appeared in the central triangular region of the plots that is typically clear of S phase cells (Fig. 4c, orange EdU+ MCM^DNA^+). We conclude that the MCM loading rate did not increase to accommodate the shorter G1 because the ↑Cyclin E1 cells had on average at least two-fold less DNA-loaded MCM in early S phase than control cells (Fig. 4d). Thus, shortening G1 length without increasing MCM loading rate causes G1 cells to enter S phase without the full complement of DNA-loaded MCM.

**Figure 4.**
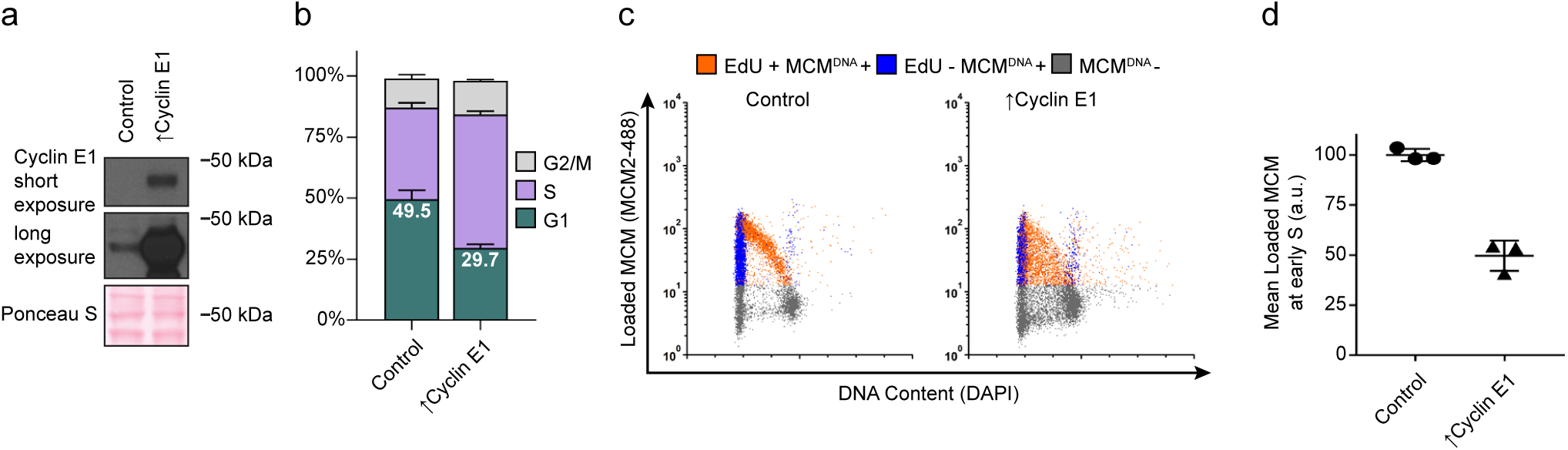
Cyclin E overproduction uncouples MCM loading and G1 length. (a) Immunoblots of a stable derivative of RPE1-hTert cells bearing an integrated doxycycline-inducible *cyclin E* construct treated with 100 ng/mL doxycycline for 72 hrs to overproduce Cyclin E1 (Cyclin E1) or with vehicle control (b) Stacked bar graphs of cell cycle distribution measured by flow cytometry for cells shown in (a); mean with error bars ± SD (n=3 biological replicates). (c) Chromatin flow cytometry of control or cyclin E-overproducing cells measuring DNA content (DAPI), DNA synthesis (EdU incorporation), and loaded MCM (anti-MCM2). (d) Mean loaded MCM in early S phase, (EdU positive, G1 DNA content) from three biological replicates; mean with error bars ± SD, unpaired two tailed t-test.***p=0.0004

### Fast loading hESCs have more Cdt1 in G1

Loading MCM complexes onto DNA requires the six subunit Origin Recognition Complex (ORC), Cdc6, and Cdt1. We hypothesized that fast MCM loading in pluripotent stem cells is achieved by increased levels of the loading proteins. To test this idea, we probed protein lysates of asynchronous cells to compare the amount of MCM loading proteins between isogenic cell lines. Total Mcm2 and ORC protein levels remained constant (Fig. 5a). The other MCM loading factors normally change in their abundance during the cell cycle due to regulated proteolysis. Cdc6 protein levels are low in G1 and high in S phase (Fig. 5b). Conversely, Cdt1 protein levels are high in G1 and low in S phase (Fig. 5b)^37,38^. Since an asynchronous population of pluripotent cells spends significantly more time in S phase than differentiated cells do, we expected Cdc6 levels to be higher in asynchronous pluripotent cells compared to their isogenic counterparts. Cdc6 was indeed higher in pluripotent cells, as was Geminin, a protein regulated in a similar manner as Cdc6 (Fig. 5a)^39^. To our surprise, even though the majority of asynchronous pluripotent cells were in S phase, a time when Cdt1 is degraded, Cdt1 levels were higher in asynchronous pluripotent cells than in isogenic differentiated cells (Fig. 5a). A similar observation was reported for mouse embryonic stem cells^40^. These data imply that Cdt1 levels are higher in G1 phase of pluripotent cells than G1 of differentiated cells, providing a potential explanation for fast MCM loading in pluripotent cells.

**Figure 5.**
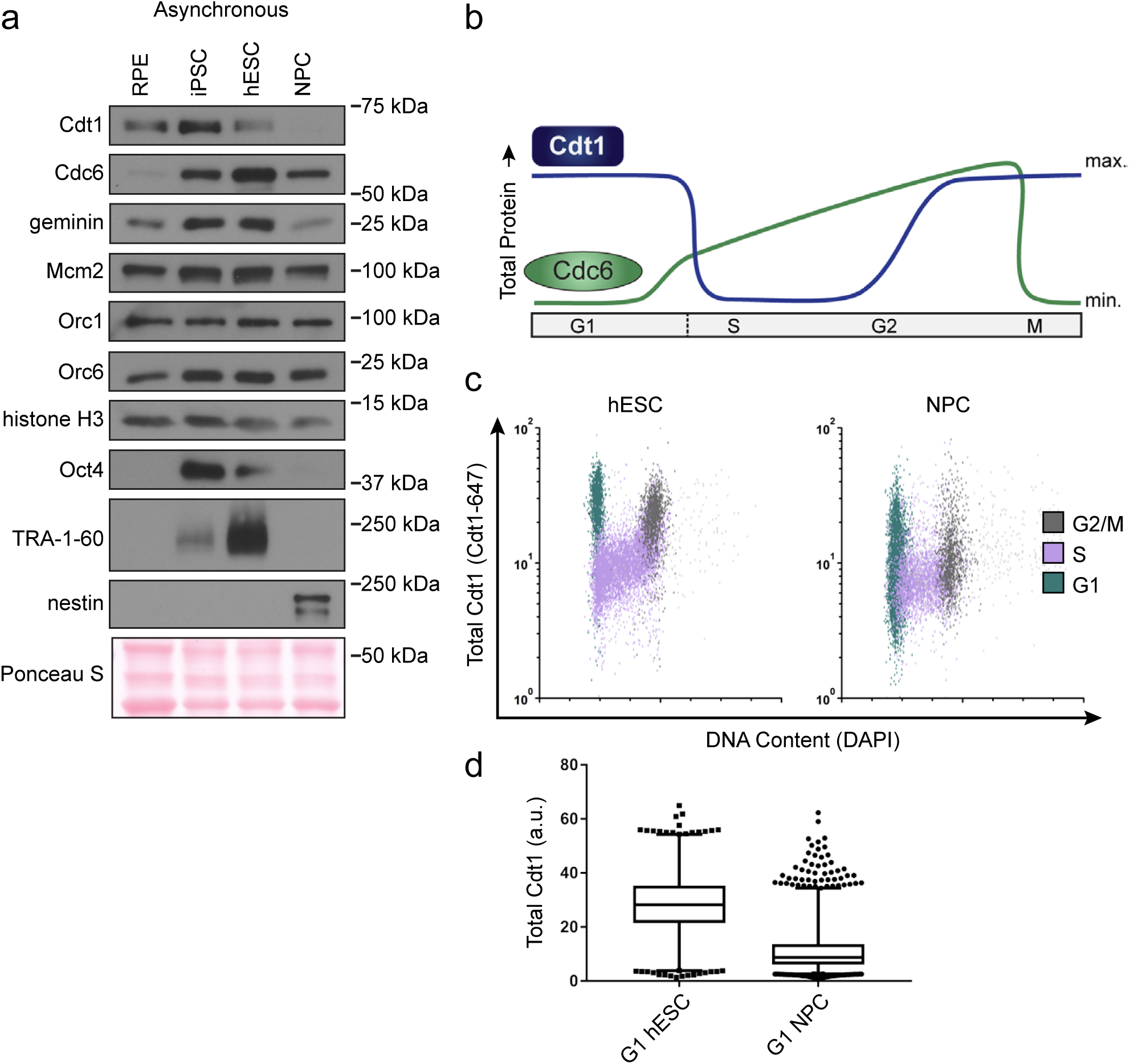
hESCs have high levels of Cdt1 in G1. (a) Immunoblots of whole cell lysates from the indicated asynchronous cell lines. (b) Expected changes in total protein levels of Cdt1 and Cdc6 during the human cell cycle. (c) Total Cdt1 detected in asynchronous cells by flow cytometry measuring DNA content (DAPI), DNA synthesis (EdU incorporation), and Cdt1 (anti-Cdt1). Green cells are G1, purple cells are S phase (EdU positive), grey cells are G2/M and >4C DNA content. (d) Box-and-whiskers plots of G1 Cdt1 concentration per cell from (C). Center line is median, outer box edges are 25^th^ and 75^th^ percentile, whiskers edges are 1^st^ and 99^th^ percentile, individual data points are lowest and highest 1%, respectively. Median G1 Cdt1 in hESCs is 2.9 fold greater, mean is 2.2 fold greater than G1 Cdt1 in NPCs, mean p=0.0504 median p=0.0243, average of 3 biological replicates.

To directly measure Cdt1 levels specifically in G1, we collected asynchronous hESCs and NPCs then fixed and stained them for Cdt1, EdU and DAPI. S phase Cdt1 degradation in hESCs is similar to differentiated cells with very low levels in S phase (purple, Fig. 5c). In contrast, the hESCs had a large population of cells with high Cdt1 levels in G1 (green cells) and significant amounts of Cdt1 in G2/M phase (grey cells), whereas the NPCs had a broad and overall lower Cdt1 distribution in G1 and very little Cdt1 in G2/M (Fig. 5d). The G1 hESCs consistently harbored significantly more Cdt1 than G1 NPCs did (2.9 fold higher median, 2.2 fold higher mean, 3 replicates (Fig. 5d)). We note that *CDT1* mRNA is modestly but consistently higher in asynchronous hESCs compared to differentiated derivatives, and that Cdt1 protein levels decrease during early differentiation coincident with the slowdown in licensing rate, but before loss of Oct4 (Fig. 3c and Supplementary Figure 4a). We postulate that the higher amount of this essential MCM loading protein specifically in G1 contributes to the fast MCM loading rate in hESCs.

### Rapid MCM loading protects hESC pluripotency

Cdt1 is essential for MCM loading, therefore reducing Cdt1 levels is expected to slow MCM loading. If MCM loading rate is linked to G1 length, then slowing MCM loading by reducing Cdt1 levels should lengthen G1. To test this prediction, we used siRNA to reduce Cdt1 in hESCs and measured changes in both MCM loading rate and G1 length (Fig. 6a, b). As expected, Cdt1 depletion reduced MCM loading rate in hESCs (Fig. 6c). Strikingly, G1 length increased coincidentally with the decrease in MCM loading rate (Fig. 6d). We obtained a complementary result by depleting the orthogonal MCM loading protein, Cdc6 (Supplementary Fig. 5a-d). A more modest Cdc6 knockdown correlated with a weaker, but detectable effect on MCM loading. Taken together, these data corroborate the close link between MCM loading rate and G1 length in hESCs.

**Figure 6.**
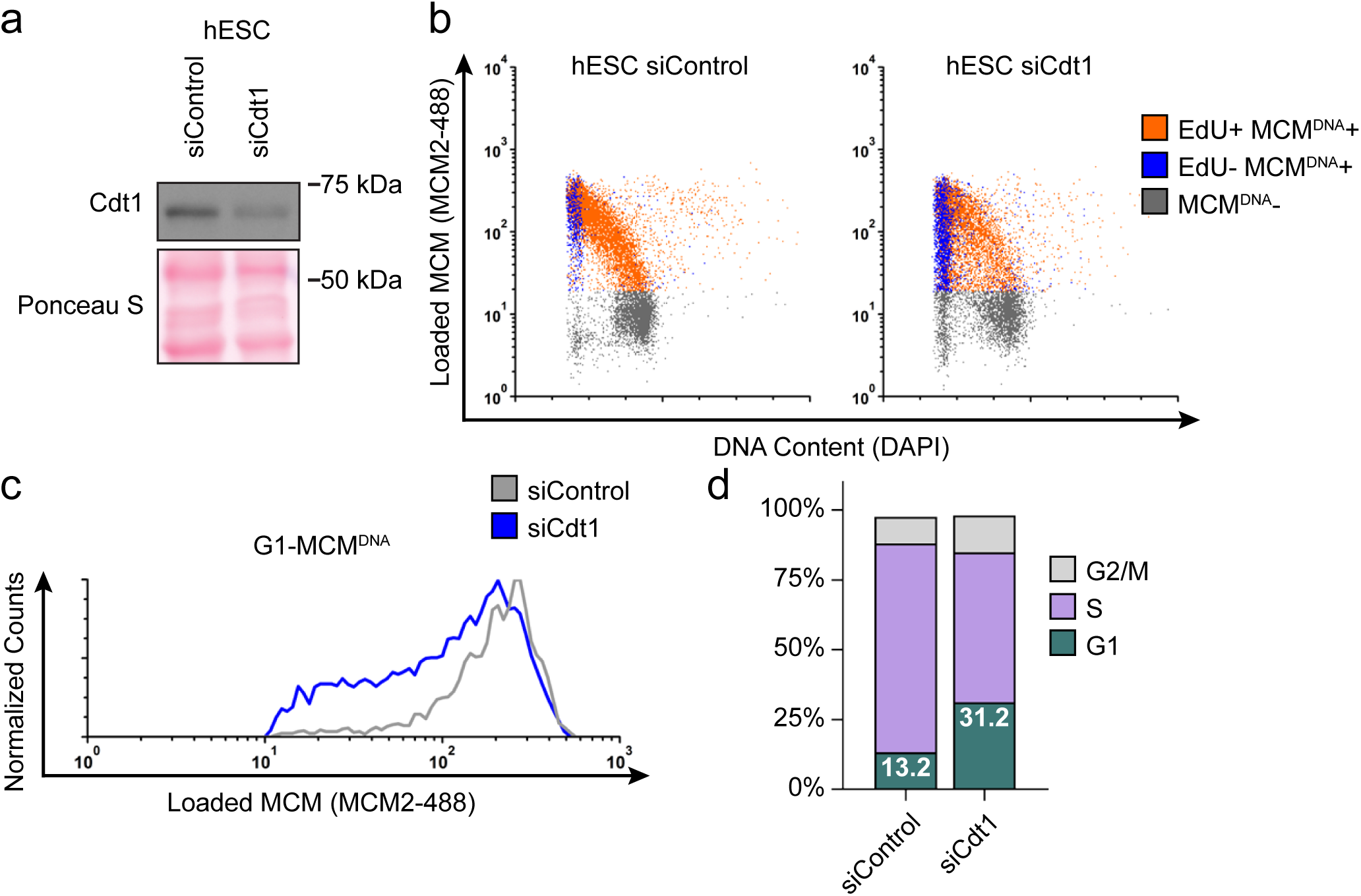
Fast MCM loading rate promotes short G1 length. (a) Immunoblot of total protein for hESCs treated with 25 nM siCdt1 or 100 nM siControl for 24 hr and labeled with EdU for 30 min prior to harvest. (b) Chromatin flow cytometry of cells from (a) as in Figure 1d. (c) Histograms of loaded MCM in G1-MCM^DNA^ cells. Counts for siCdt1 are normalized to the corresponding siControl sample. (d) Stacked bar graph of cell cycle distributions for samples in (a); representative of 2 biological replicates.

Our demonstration that slower MCM loading occurs universally during early differentiation suggested a functional link between the rate of MCM loading and pluripotency maintenance. We considered that slowing MCM loading might promote differentiation. To explore this idea, we prematurely slowed MCM loading in hESCs by Cdt1 depletion prior to inducing their differentiation (Fig. 7e). After Cdt1 depletion, we stimulated differentiation towards trophectoderm with BMP4^41^. After 48 hours, we quantified Oct4 and Cdx2 by immunostaining. (Fig. 7a). The pluripotency transcription factor Oct4 and the trophectoderm transcription factor Cdx2 reciprocally repress one another’s expression, creating a clear distinction between Oct4 positive-Cdx2 negative pluripotent cells and Oct4 negative-Cdx2 positive differentiating cells^42^. We quantified the mean fluorescence intensity of both Oct4 and Cdx2 in >18,000 cells per condition with a customized, automated CellProfiler pipeline, plotting the signal intensities for each cell in a density scatter plot (Fig. 7b, c). Stimulating control hESCs with 10 ng/mL of BMP4 slightly shifted the population towards differentiation, but most cells remained pluripotent with high Oct4 levels at this time point. Strikingly, hESCs pretreated with Cdt1 siRNA to prematurely slow MCM loading gained a substantial population of Oct4 negative-Cdx2 positive differentiating cells relative to controls that were treated similarly. To quantify the extent of differentiation, we divided the Cdx2 intensity of each cell by its Oct4 intensity, creating a single differentiation score (Fig. 7d). After 10 ng/ml BMP4 treatment, Cdt1-depleted hESCs had significantly higher scores, indicating that prematurely slowing MCM loading promoted differentiation (p<0.0001, two-tailed Mann-Whitney test). Both control cells and Cdt1-depleted cells differentiated more fully at a higher concentration of 50 ng/mL BMP4, but the Cdt1-depleted cells still differentiated further than the controls (p<0.0001, two-tailed Mann-Whitney test, Fig. 7b and data not shown). Other combinations of BMP4 concentrations or treatment times also resulted in a consistent, significant increase in differentiation in cells pretreated to slow MCM loading (p<0.0001, two-tailed Mann-Whitney test, data not shown). Importantly, the phenotype was conserved across multiple differentiation lineages, as prematurely slowing MCM loading prior to endoderm differentiation also increased the number of cells positive for the endoderm transcription factor Sox17 relative to controls at the same time point (Supplementary Fig. 6b). Additionally, slowing MCM loading by depleting a different MCM loading factor, Cdc6, also significantly promoted differentiation (p<0.0001, two-tailed Mann-Whitney test), and the more modest reduction in MCM loading (i.e. Supplementary Fig. 5b) resulted in a correspondingly weaker differentiation acceleration (Fig. S7a-e). Thus, we conclude that slow MCM loading generally promoted differentiation and by extension, that rapid MCM loading preserves pluripotency.

**Figure 7.**
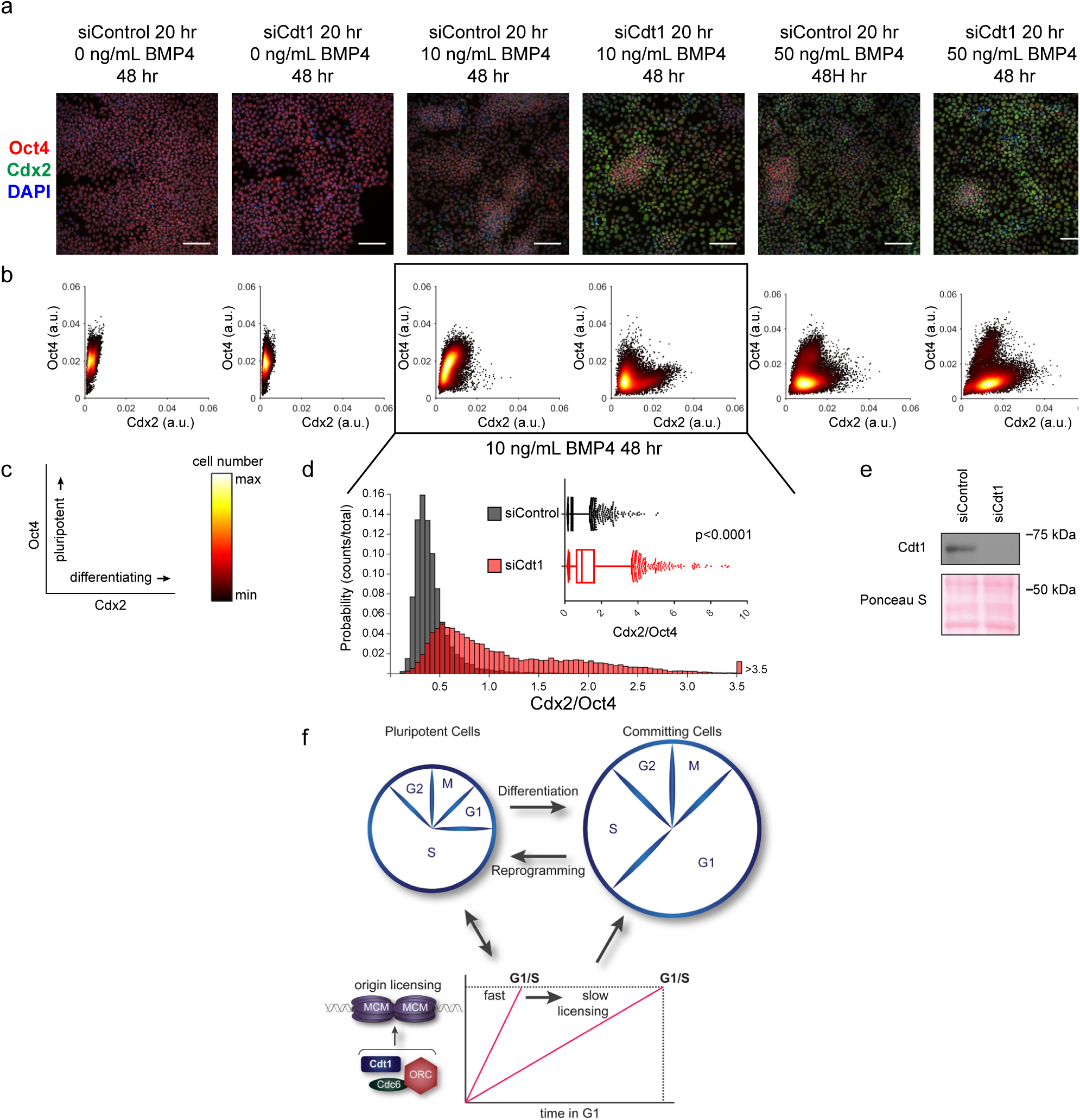
Slow MCM loading promotes differentiation. (a) Im munofluorescence microscopy of hESCs treated with 100 nM of siControl or 100 nM of siCdt1 for 20 hr and then treated with BMP4 as indicated. Cells were fixed and stained with DAPI (blue), Cdx2 antibody (green), and Oct4 antibody (red). Images are one region of 18 fields-of-view per condition; scale bar is 100 μm (see Methods). (b) Density scatterplots of mean fluorescence intensity (arbitrary units) of Oct4 and Cdx2 staining for each cell in each condition, >18,000 cells were quantified per condition. (c) Diagram of the relationship between Oct4 and Cdx2 in pluripotent and differentiated cells as plotted in (b); color bar for scatterplots in (b). (d) Histogram of mean fluorescence intensity ratio Cdx2/Oct4 for all cells in siControl and siCdt1 treated with for 10 ng/mL BMP4 for 48 hrs. Rightmost histogram bin contains all values greater than 3.5. The inset is a box-and-whiskers plot of the same data, center line is median, outer box edges are 25^th^ and 75^th^ percentile, whiskers edges are 1^st^ and 99^th^ percentile, individual data points are lowest and highest 1%, respectively. Medians are 0.3722, 0.9319, and means are 0.4285, 1.194 for siControl and siCdt1, respectively. Samples compared by two tailed Mann-Whitney test, ****p<0.0001. (e) Immunoblot for Cdt1 in whole cell lysates at 20 hr of siRNA treatment, prior to BMP4 treatment. (f) Illustration of the relationship between differentiation and MCM loading rate changes.

## DISCUSSION

In this study we demonstrate that rapid MCM loading to license replication origins is an intrinsic property of pluripotent cells. Human embryonic stem cells have a remarkably fast MCM loading rate, and reprogramming to create induced pluripotent stem cells increases MCM loading rate. Moreover, MCM loading slows concurrently with the G1 lengthening and extensive cell cycle remodeling that accompany the early stages of differentiation (Fig. 7f). To our knowledge this is the first demonstration that the rate of MCM loading is developmentally regulated. The regulated decrease in MCM loading rate is critical during differentiation, as rapid MCM loading protects pluripotency, and prematurely slowing MCM loading promotes differentiation.

Pluripotent stem cells load MCM complexes rapidly to reach similar total amounts of loaded MCM at the G1/S transition in less time than their isogenic differentiated counterparts. Although we did not detect substantial MCM loading in telophase, as suggested previously^43^, it is clear that telophase loading is an option in some cells and a requirement in cells with no detectable G1 such as *S. pombe* and in the first nuclear divisions in *D. melanogaster*^44,45^. Stem cells achieve faster MCM loading, at least in part, by particularly high Cdt1 levels in G1. These high levels are achieved not only by a modest difference in *CDT1* mRNA (Fig. 3c) but also post-transcriptionally by specific re-accumulation of Cdt1 protein in the preceding G2 phase (Fig. 5c). Cdt1 stability in G2 phase has been attributed to geminin-mediated protection from the SCF^Skp2^ E3 ubiquitin ligase in non-stem cells^46–48^. It is not clear however that geminin levels are particularly high in stem cells relative to differentiated cells (aside from differences expected from cell cycle distribution changes (Fig. 5a)^40^), so it seems unlikely that geminin drives higher Cdt1 levels in stem cells. Skp2 levels also do not change in the earliest stages of stem cell differentiation^49^. Cdt1 is protected in late S phase and G2 by cyclin A/Cdk1 activity^50^, and we thus consider it likely that the documented high CDK activity in stem cells contributes to Cdt1 stabilization in G2^51,52^. The anticipatory Cdt1 accumulation to promote MCM loading in G1 was originally proposed from experiments in cancer-derived cell lines. Our observations in stem cells suggest this strategy is employed by non-transformed cells during developmental stages that require short G1.

Other factors besides Cdt1 accumulation may also accelerate MCM loading. The hESCs we assayed have 2-3 fold greater Cdt1 protein levels in G1 relative to NPCs yet load MCM 6.5 times faster per hour than NPCs. One Cdt1 molecule can (*in vitro*) load multiple MCM complexes since Cdt1 is released into the soluble phase immediately after completing a loading reaction^53^. Stem cells may experience less Cdc6 degradation in early G1 due to nearly constitutive Cyclin E/Cdk2 activity and/or attenuated APC^Cdh1^ activity^40,52,54^. Additionally, stem cells are enriched for euchromatin, an environment that may be particularly permissive for rapid MCM loading^55^.

Rapid MCM loading may itself contribute to mechanisms that maintain short G1 phases in pluripotent cells. The origin licensing checkpoint links the amount of loaded MCM to G1 length by controlling Cdk2 activity. In that regard, overproducing Cyclin E “short-circuited” the licensing checkpoint in slow loading differentiated cells. This checkpoint has thus far only been demonstrated in p53-proficient differentiated mammalian cells^56^, but the G1 lengthening of hESCs after Cdt1 depletion suggests that pluripotent stem cells also have a functioning licensing checkpoint. Cells with fast MCM loading could satisfy this checkpoint quickly, activate Cyclin E/Cdk2, and thus spend less time in G1. Mechanisms that support short G1 length preserve pluripotency in hESCs (and promote reprogramming to iPSCs) since cells are most sensitive to differentiation cues in G1; in that regard, extending G1 phase in hESCs can increase differentiation propensity^16,20,54,57^. Recent work with quintuple knockout mice lacking all D and E type cyclins also reported that Cyclin E/Cdk2 further contributes to maintaining pluripotency by stabilizing the Oct4, Sox2, and Nanog transcription factors^58^. Fast MCM loading may have evolved as an intrinsic property of pluripotent cells to maintain high Cdk2 activity and keep G1 phase short.

Cyclin/Cdk activity is not the sole connection between the cell cycle and pluripotency. Non-CDK cell cycle-associated proteins regulate expression of key pluripotency genes including *SOX2* and *NANOG*^19,59,60^. Pluripotency transcription factors themselves regulate expression of cell cycle genes including those encoding cyclins, CDK inhibitors, and E2F3a^61–63^. On the other hand, pluripotency and cell cycle functions can be genetically uncoupled in experiments where manipulating the cell cycle didn’t alter pluripotency and vice versa^64,65^. We observe that licensing inhibition can accelerate differentiation even without greatly lengthening G1 (Supplementary Figs. 5, 7) which may point to an additional direct and cell cycle-independent link between MCM loading rate and differentiation.

We note that early differentiation is not the only setting in which rapid MCM loading during a short G1 may be relevant. Like hESCs, activated T cells have very fast cell cycles with short G1 phases^14^. Oncogenic transformation is also frequently associated with G1 shortening. It may be that the pathways linking differentiation to MCM loading rate are also coopted in some cancers to induce rapid licensing. On the other hand, a subset of cancers may proliferate in a perpetually underlicensed state that contributes to the genome instability characteristic of transformed cells. Future investigations will elucidate the molecular relationships among developmental signaling pathways, MCM loading rate, and cell cycle remodeling.

## MATERIALS AND METHODS

### Cell Culture

T98G, HEK293T, and RPE1-hTERT were cultured in Dulbecco’s Modified Eagle Medium (DMEM) supplemented with 2 mM L-glutamine and 10% fetal bovine serum (FBS) and incubated in 5% CO2 at 37 °C. ARPE-19 (male) were cultured in 1:1 DMEM:F12 supplemented with 2 mM L-glutamine and 10% fetal bovine serum and incubated in 5% CO2 at 37 °C. T98G, HEK293T, RPE1-hTERT, and ARPE-19 cells were from the ATCC and were passaged with trypsin and not allowed to reach confluency. WA09 (H9 hESCs) were cultured in mTeSR1 (StemCell Technologies) with media changes every 24 hours on Matrigel (Corning) coated dishes and incubated in 5% CO2 at 37 °C. H9s had normal diploid karyotype at passage 32 and were used from passage 32-42. ARPE-iPSCs were cultured in Essential 8 (Life Technologies) with media changes every 24 hours on Matrigel (Corning) coated dishes and incubated in 5% CO2 at 37 °C. iPSCs were used from passage 20-25. Both hESCs and iPSCs were routinely passaged every 4 days as aggregates using ReLeSR, according to manufacturer’s instructions (StemCell Technologies). The hESCs and iPSCs were only passaged as single cells in 10 μM Y-27632 2HCl (Selleck Chemicals) for experiments, as described previously ^66^. NPCs were cultured in Neural Progenitor Medium (StemCell Technologies) with media changes every 24 hours on poly-L-ornithine/Laminin (Sigma Aldrich) coated dishes and incubated in 5% CO2 at 37 °C. NPCs were passaged with StemPro Accutase (Gibco) weekly.

### Total lysate and chromatin fractionation

Cells were collected via trypsinization. For total protein lysates, cells were lysed on ice for 20 minutes in CSK buffer (300 mM sucrose, 100 mM NaCl, 3 mM MgCl_2_, 10 mM PIPES pH 7.0) with 0.5% triton x-100 and protease and phosphatase inhibitors (0.1 mM AEBSF, 1 μg/ mL pepstatin A, 1 μg/ mL leupeptin, 1 μg/ mL aprotinin, 10 μg/ ml phosvitin, 1 mM β-glycerol phosphate, 1 mM Na-orthovanadate). Cells were centrifuged at 13,000 xg at 4 °C for 5 minutes, then the supernatants were transferred to a new tube for a Bradford Assay (Biorad) using a BSA standard curve. Chromatin fractionation for immunoblotting was performed as described previously^25,26^, using CSK buffer with 1 mM ATP, 5 mM CaCl_2_, 0.5% triton x-100 and protease and phosphatase inhibitors to isolate insoluble proteins and S7 nuclease (Roche) to release DNA bound proteins. A Bradford Assay (Biorad) was performed for equal loading.

### Immunoblotting

Samples were diluted with SDS loading buffer (final: 1% SDS, 2.5% 2-mercaptoethanol, 0.1% bromophenol blue, 50 mM Tris pH 6.8, 10% glycerol) and boiled. Samples were run on SDS-PAGE gels, then the proteins transferred onto polyvinylidene difluoride membranes (Thermo Fisher). Membranes were blocked at room temperature for 1 hour in either 5% milk or 5% BSA in Tris-Buffered-Saline-0.1%-tween-20 (TBST). After blocking, membranes were incubated in primary antibody overnight at 4 °C in either 1.25% milk or 5% BSA in TBST with 0.01% sodium azide. Blots were washed with TBST then incubated in HRP-conjugated secondary antibody in either 2.5% milk or 5% BSA in TBST for 1 hour, washed with TBST, and then membranes were incubated with ECL Prime (Amersham) and exposed to autoradiography film (Denville). Equal protein loading was verified by Ponceau S staining (Sigma Aldrich). Antibodies used for immunoblotting were: Mcm2, (BD Biosciences, Cat#610700), Mcm3, (Bethyl Laboratories, Cat#A300-192A), Cdt1, (Cell Signaling Technologies, Cat#8064S), Cdc6, (Santa Cruz Biotechnology, Cat#sc-9964), Oct4, (Abcam, Cat#ab19857), Cdx2, (Abcam, Cat#ab76541), Cyclin E1, (Santa Cruz Biotechnology, Cat#sc-198), Orc1, (Bethyl Laboratories, Cat#A301-892A), Orc6, (Santa Cruz Biotechnology, Cat#sc-32735), geminin, (Santa Cruz Biotechnology, Cat#sc-13015), Histone H3, (Cell Signaling Technologies, Cat#4499S), TRA-1-60, (Invitrogen, Cat#41-1000), nestin, (Abcam, Cat#ab22035).

### Flow Cytometry

For EdU labeled samples, cells were incubated with 10 uM EdU (Santa Cruz Biotechnology) for 30 minutes prior to collection. For total protein flow cytometry, cells were collected with trypsin and resuspended as single cells, washed with PBS, and fixed with 4% paraformaldehyde (Electron Microscopy Sciences) in PBS for 15 minutes at room temperature, then 1% BSA-PBS was added, mixed and cells were centrifuged at 1000 xg for 7 minutes (and for all following centrifuge steps) then washed with 1% BSA-PBS and centrifuged. Fixed cells were permeabalized with 0.5% triton x-100 in 1% BSA-PBS at room temperature for 15 minutes, centrifuged, then washed once with 1% BSA, PBS and centrifuged again before labeling. For chromatin flow cytometry, cells were collected with trypsin and resuspended as single cells, washed with PBS, and then lysed on ice for 5 minutes in CSK buffer with 0.5% triton x-100 with protease and phosphatase inhibitors. Next, 1% BSA-PBS was added and mixed, then cells were centrifuged for 3 minutes at 1000 xg, then fixed in 4% paraformaldehyde in PBS for 15 minutes at room temperature. Next, 1% BSA-PBS was added, mixed and cells were centrifuged then washed again before labeling. The labeling methods for total protein samples and chromatin samples were identical. For DNA synthesis (EdU), samples were centrifuged and incubated in PBS with 1 mM CuSO_4_, 1 μM fluorophore-azide, and 100 mM ascorbic acid (fresh) for 30 minutes at room temperature in the dark. 1% BSA-PBS + 0.1% NP-40 was added, mixed and centrifuged. Samples were resuspended in primary antibody in 1% BSA-PBS + 0.1% NP-40 and incubated at 37 °C for 1 hour in the dark. Next, 1% BSA-PBS + 0.1% NP-40 was added, mixed and centrifuged. Samples were resuspended in secondary antibody in 1% BSA-PBS + 0.1% NP-40 and incubated at 37 °C for 1 hour in the dark. Next, 1% BSA-PBS + 0.1% NP-40 was added, mixed and centrifuged. Finally, cells were resuspended in 1% BSA-PBS + 0.1% NP-40 with 1 μg/mL DAPI (Life Technologies) and 100 μg/mL RNAse A (Sigma Aldrich) and incubated overnight at 4 °C in the dark. Samples were run on a CyAn ADP flow cytometer (Beckman Coulter) and analyzed with FCS Express 6 software (De Novo). The following antibody/fluorophore combinations were used: (1): Alexa 647-azide (Life Technologies), primary: Mcm2 (BD Biosciences, Cat#610700), secondary: Donkey anti-mouse-Alexa 488 (Jackson ImmunoResearch), DAPI. (2): Alexa 488-azide (Life Technologies), primary: Cdt1 (Abcam, Cat#610700), secondary: Donkey anti-rabbit-Alexa 647 (Jackson ImmunoResearch), DAPI. Cells were gated on FS-area vs SS-area. Singlets were gated on DAPI area vs DAPI height. The positive/negative gates for EdU and MCM were gated on a negative control sample, which was treated with neither EdU nor primary antibody, but incubated with 647-azide and the secondary antibody Donkey anti-mouse-Alex 488 and DAPI to account for background staining (Supplementary Figure 1).

### Doubling Time

Doubling time was calculated by plating equal number of cells as described above and counting cell number over time using a Luna II automated cell counter (Logos Biosystems) at 24, 48, and 72 hours after plating. Three or four wells were counted as technical replicates at each timepoint. GraphPad Prism’s regression analysis was used to compute doubling time, and multiple biological replicates were averaged for a final mean doubling time. ARPE-19s were counted 4 times, hESCs and NPCs 3 times, and iPSCs 2 times.

### Cell Synchronization

To synchronize cells in G1, T98G cells were grown to 100% confluency, washed with PBS, and incubated for 72 hr in 0.1% FBS, DMEM, L-glutamine. After serum-starvation, cells were re-stimulated by passaging 1:3 with trypsin to new dishes in 20% FBS, DMEM, L-glutamine, collecting cells 10 hr and 12 hr post-stimulation. To synchronize T98G cells in early S phase, cells were treated as for G1, except 1 mM Hydroxyurea (Alfa Aesar) was added to the media upon re-stimulating and cells were collected 18 hr post-stimulation. To synchronize cells in mid-late S, cells were treated as in early S, then at 18 hr post-stimulation cells were washed with PBS and released into 10% FBS, DMEM, L-Glutamine, collecting 6 hr, 8 hr post release.

### Cloning

The pInducer20-Cyclin E plasmid was constructed using the Gateway cloning method (Invitrogen). The *att*B sites were added to Cyclin E1 cDNA by PCR using Rc/CMV cyclin E plasmid as a template and BP-cycE-F (5′ GGGGACAAGTTTGTACAAAAAAGCAGGCTACCATGAAGGAGGACGGCGGC) and BP-cycE-R primers (5′ GGGGACCACTTTGTACAAGAAAGCTGGGTTCACGCCATTTCCGGCCCGCT) ^67^. The PCR product was recombined with pDONR221 plasmid using BP clonase (Invitrogen) according to the manufacturer’s instructions and transformed into DH5α to create pENTR221-Cyclin E1. Then the LR reaction was performed between pInducer20 and pENTR221-Cyclin E1 using LR Clonase (Invitrogen) according to manufacturer’s instructions and transformed into DH5α to create pInducer20-Cyclin E1.

### Inducible Cyclin E

To package lentivirus, pInducer20-Cyclin E1 was co-transfected with ΔNRF and VSVG plasmids into HEK293T using 50 μg/mL Polyethylenimine-Max (Aldrich Chemistry). Viral supernatant was transduced with 8 ug/mL Polybrene (Millipore) onto RPE1-hTERT cells overnight. Cells were selected with 500 ug/mL neomycin (Gibco) for 1 week. To overproduce Cyclin E1, cells were treated with 100 ng/mL doxycycline (CalBiochem) for 72 hr in 10% FBS, DMEM, L-glutamine. Control cells were the Inducer20-Cyclin E1 without doxycycline.

### siRNA transfections

For siRNA treatment, Dharmafect 1 (Dharmacon) was mixed in mTeSR1 with the appropriate siRNA according to the manufacturer’s instructions, then diluted with mTeSR1 and added to cells after aspirating old media. The final siRNA concentrations were: 100 nM siControl (Luciferase), 25 or 100 nM siCdt1, or a mixture of two siCdc6 (2144 and 2534 at 50 nM each). The Cdt1 siRNA mix was incubated on cells for either 20 or 24 hours, then changed to new mTeSR1 without siRNA. The Cdc6 siRNA mix was incubated on cells for 24 hours, then changed to new mTeSR1 without siRNA for 8 hours (32 total hours). The siRNA were described previously ^68,69^.

### Differentiation

Trophectoderm: hESCs were passaged as single cells at 7x10^3^ /cm^2^ in mTeSR1 with 10 μM Y-27632 2HCl onto Matrigel-coated plates. 24 hours later, the media was changed to start differentiation with fresh mTeSR1 with 100 ng/mL BMP4 (R&D Systems), and 24 hours later the media was changed to fresh mTeSR1 with 100 ng/mL BMP4 for 48 total hours of differentiation.

Neuroectoderm: hESCs were differentiated using a monolayer-based protocol in Neural Induction Medium: hESCs were passaged as single cells at 5.2x10^4^ /cm^2^ in STEMdiff™ Neural Induction Medium (StemCell Technologies) with 10 μM Y-27632 2HCl onto Matrigel coated plates, and plating started the differentiation. 24 hours later, the media was changed to fresh Neural Induction Medium for another 24 hours for 48 hours total differentiation. To derive NPCs, hESCs were differentiated in Neural Induction medium (StemCell Technologies) using the Embryoid Body Neural Induction protocol according to manufacturer’s instructions, similar to previous reports ^70^. Once generated, NPCs were maintained in Neural Progenitor Medium (StemCell Technologies).

Mesoderm: hESCs were passaged as single cells at 3x10^4^/cm^2^ in mTeSR1 with 10 μM Y-27632 2HCl onto Matrigel coated plates. 24 hours later, the media was changed to start differentiation. Cells were washed with Advanced RPMI 1640 (Gibco), then incubated in Advanced RPMI 1640 with B27 minus insulin (Gibco), 2 mM L-glutamine, and 8 μM CHIR-99021 (Selleck Chemicals). At 24 hours after changing the media, cells were washed with Advanced RPMI 1640 then incubated in Advanced RPMI 1640 with B27 minus insulin (Gibco), 2 mM L-glutamine, without CHIR-99021 for 24 hours for a total of 48 hours of differentiation.

Endoderm: hESCs were passaged as single cells at 4x10^3^/cm^2^ in mTeSR1 with 10 μM Y-27632 2HCl onto Matrigel-coated plates. The next day the media was changed to fresh mTeSR1 without Y-27632 2HCl. 24 hours later, the media was changed to start differentiation. The cells were washed with Advanced RPMI 1640 (Gibco), then incubated in Advanced RPMI 1640 with 0.2% FBS, 2 mM L-glutamine, 100 ng/mL Activin A (R&D Systems) and 2.5 μM CHIR-99021. At 24 hours after changing the media, cells were washed with Advanced RPMI 1640 then incubated in Advanced RPMI 1640 with 0.2% FBS, 2 mM L-glutamine, 100 ng/mL Activin A, without CHIR-99021 for 24 hours for a total of 48 hours of differentiation.

### Phase contrast microscopy

Phase contrast images were acquired with an Axiovert 40 CFL inverted microscope, 20x objective.

### Immunofluorescence Microscopy

For immunofluorescence microscopy, hESCs were plated as single cells in mTeSR1 with 10 μM Y-27632 2HCl in Matrigel-coated, 24 well, #1.5 glass bottom plates (Cellvis) at 7x10^3^/cm^2^ for siCdt1, Trophectoderm, at 5x10^3^ for siCdc6, Trophectoderm, and at 4x10^3^/cm^2^ for siCdt1, Endoderm. Cells were incubated with siCdt1 for 20 hours (Trophectoderm), 24 hours (Endoderm) or siCdc6 for 32 hours (Trophectoderm) all in parallel with siControl as described above (siRNA transfections). After siRNA treatment, cells were differentiated as described above (Differentiation) with the following modifications: For trophectoderm, multiple BMP4 concentrations and treatment times were used as indicated (Fig. 7, Supplementary Fig. 6). For treatment less than 48 hours, cells were incubated in mTeSR1 after siRNA treatment until starting differentiation. (Example: 12 hours of mTeSR1 then 36 hours of BMP4, for a total of 48 hours). For endoderm, the first RPMI/Activin/CHIR-99021 was immediately after siRNA, without a day of incubation in mTeSR1. After differentiation, cells were fixed in 4% paraformaldehyde in PBS for 15 minutes at room temperature, washed with PBS, and permeabalized with 5% BSA, PBS, 0.3% triton x-100 at 4 °C overnight. Next, cells were incubated in primary antibody in 5% BSA, PBS, 0.3% triton x-100 at 4 °C overnight. Cells were washed with PBS at room temperature, then incubated in secondary antibody in 5% BSA, PBS, 0.3% triton x-100 at room temperature for 1 hour. Cells were washed with PBS, then incubated in 1 μg/mL DAPI in PBS for 10 minutes at room temperature, then washed with PBS. For trophectoderm the primary antibodies were Oct4 (Millipore, Cat#MABD76) and Cdx2 rabbit (Abcam, Cat#ab76541), the secondary antibodies were goat anti-mouse-Alexa 594, donkey anti-rabbit-Alexa 488. For endoderm the primary antibody was Sox17 (R&D Systems, Cat#AF1924), the secondary antibody was donkey anti-goat-Alexa 594 (Jackson ImmunoResearch). Cells were imaged in PBS on a Nikon Ti Eclipse inverted microscope with an Andor Zyla 4.2 sCMOS detector. Images were taken as 3x3 scan of 20x fields with a 0.75 NA objective, stitched with 15% overlap between fields using NIS-Elements Advanced Research Software (Nikon). Shading correction was applied within the NIS-Elements software before acquiring images. Raw images were quantified using a custom CellProfiler pipeline.

### qPCR (Figure 3)

RNA lysates were prepared using Norgen Biotek’s Total RNA Purification Kit (Cat. 37500). Lysates were first treated with Promega RQ1 RNase-Free DNase (Promega), and then converted to cDNA using Applied Biosystem’s High-Capacity RNA-to-cDNA Kit (Cat. 4387406). Quantitative real-time PCR (qPCR) with SYBR Green (Bio-Rad; SsoAdvanced Universal SYBR Green Supermix, Cat. 1725271) was carried out to assess gene expression. All results were normalized to *ACTB*. Primers for qPCR were ordered from Eton Bioscience. Primers:

POU5F1-F: 5′-CCTGAAGCAGAAGAGGATCACC,

POU5F1-R 5′-AAAGCGGCAGATGGTCGTTTGG,

CDX2-F 5′-ACAGTCGCTACATCACCATCCG,

CDX2-R 5′-CCTCTCCTTTGCTCTGCGGTTC,

T-F 5′-CTTCAGCAAAGTCAAGCTCACC,

T-R 5′-TGAACTGGGTCTCAGGGAAGCA,

SOX17-F 5′-ACGCTTTCATGGTGTGGGCTAAG,

SOX17-R 5′-GTCAGCGCCTTCCACGACTTG,

CDT1-F 5′-GGAGGTCAGATTACCAGCTCAC,

CDT1-R, 5′-TTGACGTGCTCCACCAGCTTCT,

SOX2-F 5′-CTACAGCATGATGCAGGACCA,

SOX2 -R 5′-TCTGCGAGCTGGTCATGGAGT,

PAX6-F 5′-AATCAGAGAAGACAGGCCA,

PAX6-R 5′-GTGTAGGTATCATAACTC,

ACTB-F 5′-CACCATTGGCAATGAGCGGTTC,

ACTB-R 5′-AGGTCTTTGCGGATGTCCACGT.

### Generating ARPE-19-iPS cells

ARPE-19-iPS cells were derived from the human retinal pigment epithelial cell line ARPE-19 by reprogramming with CytoTune-iPS 2.0 Sendai reprogramming kit (Invitrogen) following the manufacturer’s instructions.

Briefly, two days before Sendai virus transduction, 100,000 ARPE-19 cells were plated into one well of a 6-well plate with ATCC-formulated DMEM:F12 medium and were transduced with the CytoTune™ 2.0 Sendai reprogramming vectors at the MOI recommended by the manufacturer 48 hours later (d0). The medium was replaced with fresh medium every other day starting from one day after transduction (d1). At day 7, transduced cells were replated on Matrigel coated 6-well plates. Cells were fed with Essential 8 medium every day. Colonies started to form in 2-3 weeks and were ready for transfer after an additional week. Undifferentiated colonies were manually picked and transferred to Matrigel coated 6-well plates for expansion. After two rounds of subcloning and expansion (after passage 10), RT-PCR was used to verify whether iPS cells were vector-free with the primer sequences published in the manufacturer’s manual.

After iPS cells became virus-free, they were submitted to the University of Minnesota Cytogenomic Laboratory for karyotype analysis. This analysis indicated that the ARPE-19-iPS cells have normal karyotypes.

### Immunofluorescence characterization of ARPE-19-iPS cells

To examine pluripotency markers, iPS cells were fixed with 4% paraformaldehyde for 20 minutes. If nuclear permeation was required, cells were treated with 0.2% triton-x-100 in phosphate-buffered saline (PBS) for 30 minutes, blocked in 3% bovine serum albumin in PBS for 2 hours, and incubated with the primary antibody overnight at 4°C. Antibodies targeting the following antigens were used: TRA1-60 (MAB4360, 1:400), TRA1-81 (MAB4381, 1:400), stage-specific embryonic antigen-4 (MAB4304, 1:200), and stage-specific embryonic antigen-3 (MAB-4303, 1:200), all from Millipore/Chemicon (Billerica, MA), OCT3/4 (AB27985, 1:200) from Abcam (Cambridge, MA), and NANOG (EB068601:100) from Everest (Upper Heyford, Oxfordshire, UK). Cells were incubated with secondary Alexa Fluor Series antibodies (all 1:500, Invitrogen) for 1 hour at room temperature and then with DAPI for 10 minutes. Images were examined using an Olympus FluoView 1000m IX81 inverted confocal microscope and analyzed with Adobe Photoshop CS6. Direct alkaline phosphatase (AP) activity was analyzed as per the manufacturer’s recommendations (Millipore).

### Bisulfite sequencing and methylation analysis

Genomic DNA was isolated using the DNeasy Blood and Tissue kit (Qiagen) per manufacturer’s recommendations for isolation from mammalian cells. Bisulfite conversion was performed using the Epitect Bisulfite kit (Qiagen) according to the manufacturer’s protocol for low amounts of DNA. Single-step PCR amplification of the NANOG and OCT4 promoter regions were conducted using Accuprime Supermix II (Invitrogen). Amplification products were visualized by gel electrophoresis and bands were excised and purified using the QIAquick Gel Extraction kit (Qiagen). Purified PCR products were inserted into the PCR4-TOPO vector (Invitrogen) and individual clones were sequenced. Alignment and methylation analysis were performed using the online QUMA program (http://quma.cdb.riken.jp/). Sequenced clones with at least 90% non-CpG cytosine conversion and at least 90% sequence homology were retained for analysis.

### Quantitative reverse transcriptase PCR (Supplementary Figure 2)

RNA was isolated using RNeasy Mini kit (Qiagen) and treated with TURBO DNA-free (Ambion, Austin, TX). First-strand cDNA was synthesized using a Superscript III First-Strand Synthesis SuperMix (Invitrogen). Reverse transcriptase-PCR was performed using TaqMan Gene Expression Assays and TaqMan Universal PCR Master Mix, No AmpErase UNG (Applied Biosystems, Carslbad, CA) as per the manufacturer’s protocol.

TaqMan gene expression assays used were OCT4 (Hs04260367-gH), SOX2 (Hs01053049-sl), NANOG (Hs04399610-g1), KLF4 (Hs00358836-m1), MYC (Hs00153408-m1), LIN28 (Hs00702808-s1), REXO1 (Hs00810654-m1), ABCG2 (Hs1053790-m1), DNMT3 (Hs00171876-m1), with GAPDH (Hs99999905-m1) used as an endogenous control. Expression levels were measured in duplicate. For genes with expression below the fluorescence threshold, the cycle threshold (Ct) was set at 40 to calculate the relative expression. Analysis was performed using an ABI PRISM 7500 sequence detection system (Applied Biosystems).

### Teratoma analysis

ARPE-19-iPS cells contained in a mixture of DMEM/F12, Matrigel and collagen were implanted onto the hind flank of NSG mice (n=5) until a palpable mass formed. Teratoma tissue was excised for histological examination following embedding and staining by haematoxylin and eosin. Experiments were conducted with the approval of the Institutional Animal Care and Research Committee at the University of Minnesota.

### Ergodic Rate Analysis

Ergodic Rate Analysis for cell cycle data was based on previously published work ^33^. First, the raw data from flow cytometry files were extracted using FCSExtract Utility (Earl F Glynn) to comma separated value (.csv) files. The data in the .csv files were then gated in FCS Express 6 (De Novo Software) and the data for only G1-MCM^DNA^ were exported. MCM negative cells were excluded based on the negative control sample (see Flow Cytometry). The mean MCM loading rate was calculated in MATLAB (MathWorks). To calculate the mean MCM loading rate, the G1-MCM^DNA^ were subdivided into 10 equal sized bins, with rate calculated for each bin, and all 10 rates were averaged together for a mean MCM loading rate. The rate calculation was based on the formula from Kafri et al:

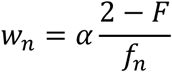

w_n_ = MCM loading rate in bin n

α = ln(2) / doubling time (Supplementary Figure 2)

F = number of G1-MCM^DNA^ cells / total number of cells in sample. F was calculated from FCS Express and entered into MATLAB manually (Supplementary Figure 2).

f_n_= number of cells in bin n / number of G1-MCM^DNA^ cells

The bins were created in MATLAB (Figure S2). To control for small day to day differences in raw data from staining intensities, the histogram edges were defined with the first bin starting at the lowest MCM value and the last bin ending at the highest MCM value, divided into 10 equal sized bins between the lowest and highest MCM value. The 10 w^n^ were then averaged for a final mean w per sample. Sample MATLAB code:

~~~
alpha_iPSC=log(2)/15.64;
F_iPSC_1=0.0801;
%calculate lowest and highest MCM values%
maxMCM_iPSC_1=max(iPSC_1(:,1));
minMCM_iPSC_1=min(iPSC_1(:,1));
%create histogram with 10 bins and specified first and last bin limits%
h10_iPSC_1=histogram(iPSC_1(:,1), ‘NumBins’, 10, ‘BinLimits’, [minMCM_iPSC_1, maxMCM_iPSC_1]);
%calculate fn within each bin%
totalf_iPSC_1=size(iPSC_1);
fn10_iPSC_1=(h10_iPSC_1.Values)/totalf_iPSC_1(1,1);
%calculate mean w%
w10mean_iPSC_1=mean(alpha_iPSC .*(2-F_iPSC_1)./fn10_iPSC_1);
~~~

The mean MCM loading rate was calculated for 3 biological replicates for each cell line, and the replicates were averaged using GraphPad Prism for further statistical analysis. We cannot use ergodic rate analysis on actively differentiating cells (e.g. Figure 3) because they are not at steady state.

### Quantification and Statistical Analysis

Statistical analysis was performed with GraphPad Prism 7 using unpaired, two-tailed t test (displayed as mean ±SD) or two-tailed Mann-Whitney test as indicated in figure legends. Significance levels were set at ^*^p≤ 0.05, ^**^p≤ 0.01, ^***^p≤ 0.001, ^****^p≤ 0.0001. All experiments were performed a minimum of 2 times, and representative data are shown in figures.

## AUTHOR CONTRIBUTIONS

J.P.M. designed and performed the experiments and analyzed the results. R.D. and P.C. performed experiments. R.M.B. W.C, K.T., B.R.W., J.T. and A.K.B. generated, validated the iPSCs and associated data in Supplementary Figure 2. J.E.P. and J.G.C. designed experiments and supervised the project. J.P.M and J.G.C. wrote the manuscript with input from the other authors.

## ACKNOWLEDGMENTS

We are grateful to Ran Kafri for advice on ergodic rate analysis. We thank Jeffrey Jones for managerial assistance, Dr. Sam Wolff for microscopy assistance, the UNC Human Pluripotent Stem Cell Core stem cell culture assistance and A. Adams, S. Wong, M. Consuegra, S. Goraya, and S.Sisk for technical assistance. We also thank Dr. Robert Duronio, Dr. Michael Emanuele, and the Cook lab for helpful discussions. We thank Ron McElmurry and Megan Riddle at the University of Minnesota for conducting teratoma assays, and the Cytogenomics Shared Resource (P30 CA077598) at the University of Minnesota Masonic Cancer Center for karyotype analysis. The UNC Flow Cytometry Core Facility is supported in part by P30 CA016086. This work was supported by a fellowship from the NSF (DGE-1144081) to J.P.M and by grants from the NIH to J.G.C. (GM083024 and GM102413), R.M.B. (T32-CA009138) J.E.P. (DP2-HD091800) and A.K.B. (GM074917). Additional funding was provided by the W.M. Keck foundation to J.E.P. and J.G.C; J.T. is supported by the Tulloch Chair in Stem Cell Biology, Genetics and Genomics at the University of Minnesota.

## COMPETING INTERESTS

We have no competing financial or non-financial interests.

## SUPPLEMENTAL FIGURES

**Supplementary Figure 1.**
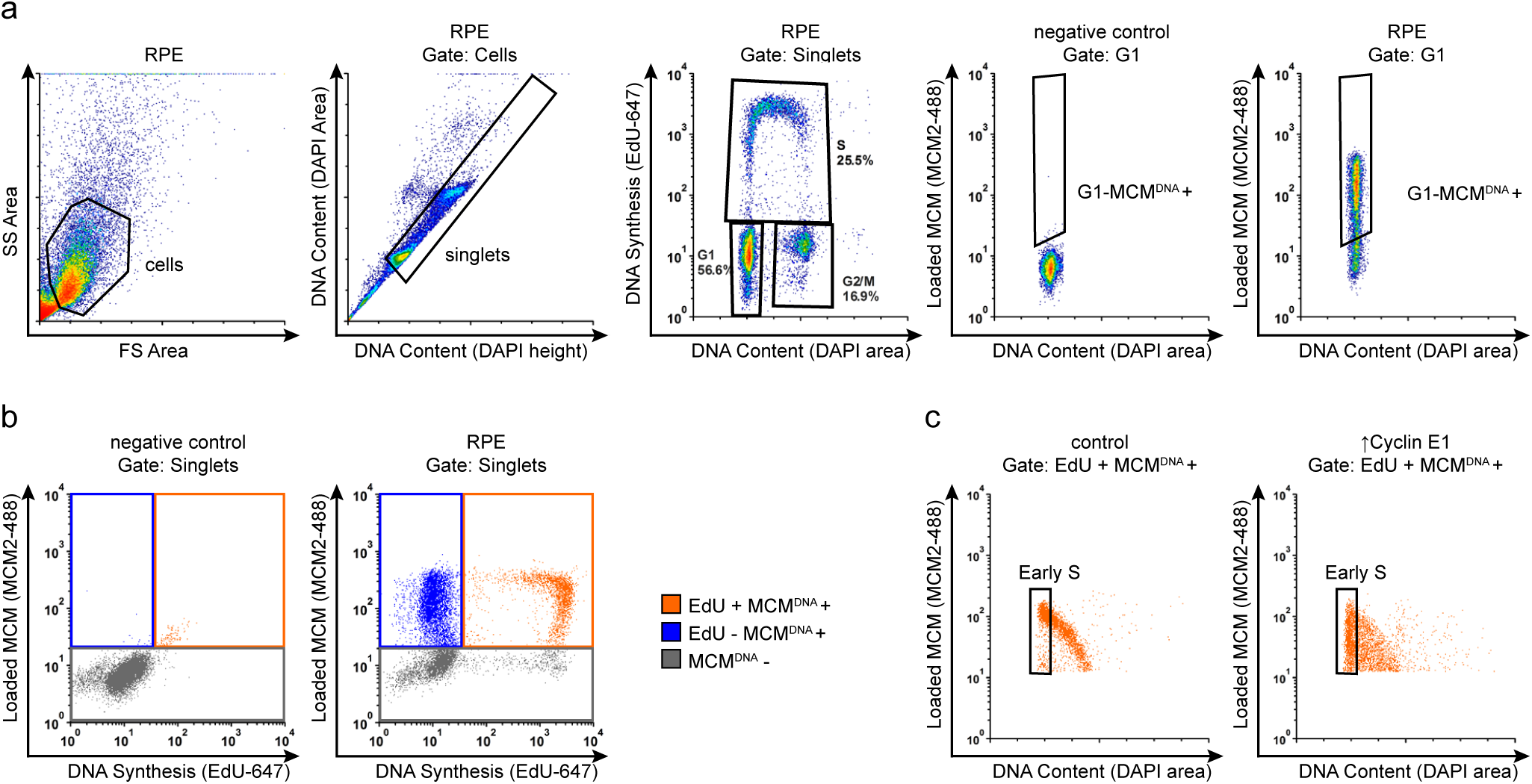
Flow Cytometry Gating. (a) Example flow cytometry gating with chromatin extracted ARPE-19 (RPE) cells from Figure 1d. Gating for cells on FS-area vs SS-area, singlets on DAPI-height vs DAPI area, cell cycle phases on DAPI-area vs EdU-647, and G1-MCM^DNA^+ on DAPI-Area vs MCM2-488. (b) Example flow cytometry gating with chromatin extracted ARPE-19 cells from Figure 1d. (left) Negative control to define background without EdU labeling or MCM2 primary antibody. Cells were stained with DAPI, subjected to EdU detection chemistry, and stained with secondary antibody. (right) Chromatin flow cytometry of ARPE-19 measuring DNA content (DAPI), DNA synthesis (EdU-647), and loaded MCM (anti-MCM2). Gates define color schemes for color dot plots. (c) Gating for early S phase for cells in Figure 4d, EdU +, MCM^DNA^+, G1 DNA Content.

**Supplementary Figure 2.**
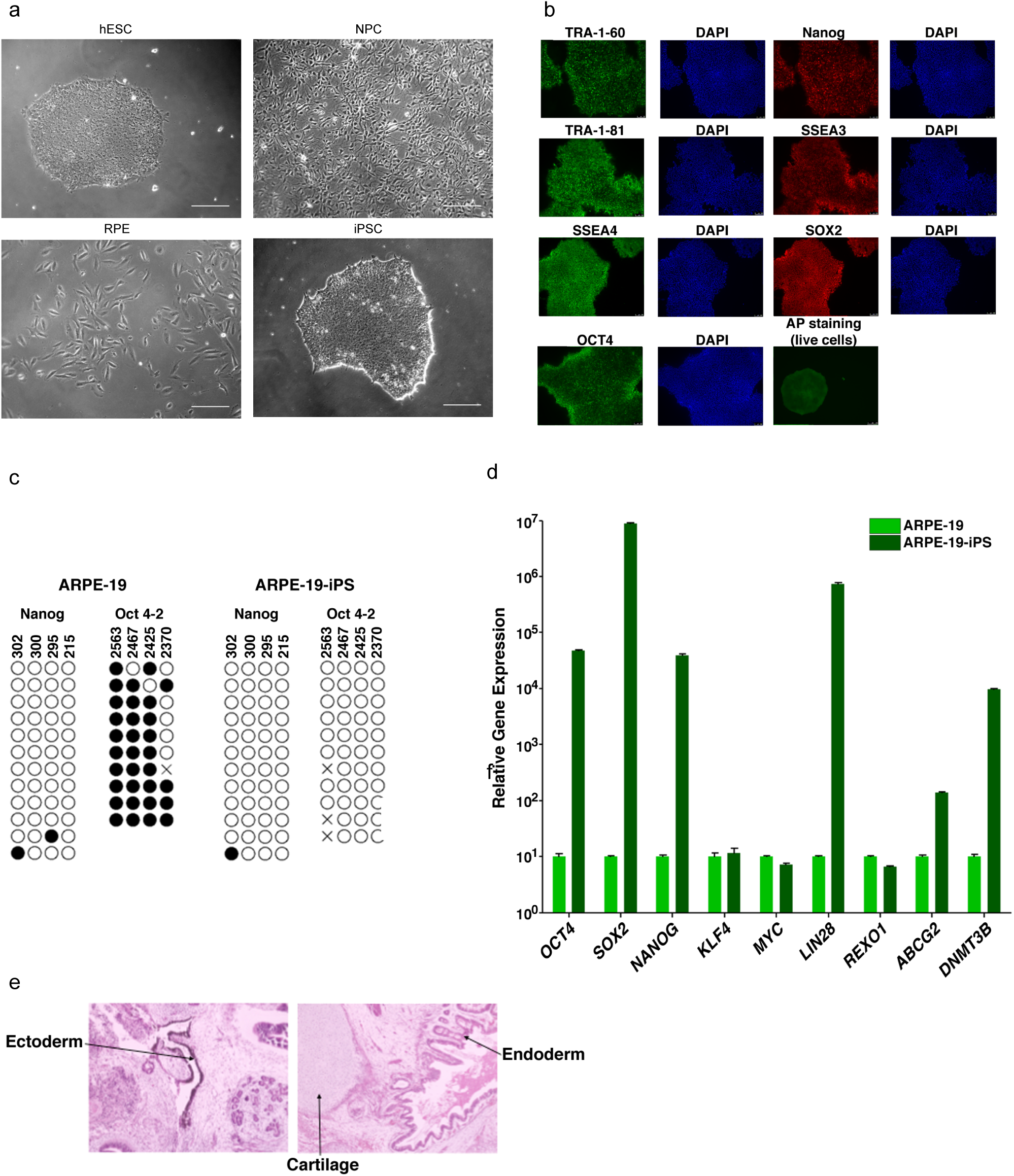
Characterization of pluripotent and differentiated cells. (a) Representative phase contrast images in greyscale of indicated cell lines from Figure 1d. Scale bar is 50 μm. (b) Immunofluorescence analysis of iPS cells visualizing TRA-1-60, TRA-1-81, SSEA or OCT4 (in green) and Nanog, SSEA3 or SOX2 (in red). DAPI counterstaining for DNA indicates the nuclei of individual cells in each colony. Live alkaline phosphatase (AP) staining was also performed as a positive indicator of stem cells. (c) Bisulfite sequencing analysis of the NANOG and OCT4-2 promoter regions in ARPE-19 and ARPE-19-iPS cells is indicated. Methylated CpGs are indicated with a closed circle, and unmethylated CpGs are indicated with an open circle. An ‘X’ indicates a mismatch or gap in the bisulfite sequence. The CpG position relative to the downstream transcription start site is shown above each row. (d) Quantitative RT-PCR analysis showing the relative gene expression of OCT4, SOX2, NANOG, KLF4, MYC, LIN28, REXO1, ABCG2 and DNMT3B in ARPE-19 and ARPE-19-iPS cells (normalized to GAPDH). Error bars indicate standard error. (e) Hematoxylin and eosin staining of teratoma sections from immunodeficient mice injected with ARPE-19-iPS cells show cartilage, as well as endoderm and ectoderm teratoma formed by ARPE-19-iPS.

**Supplementary Figure 3.**
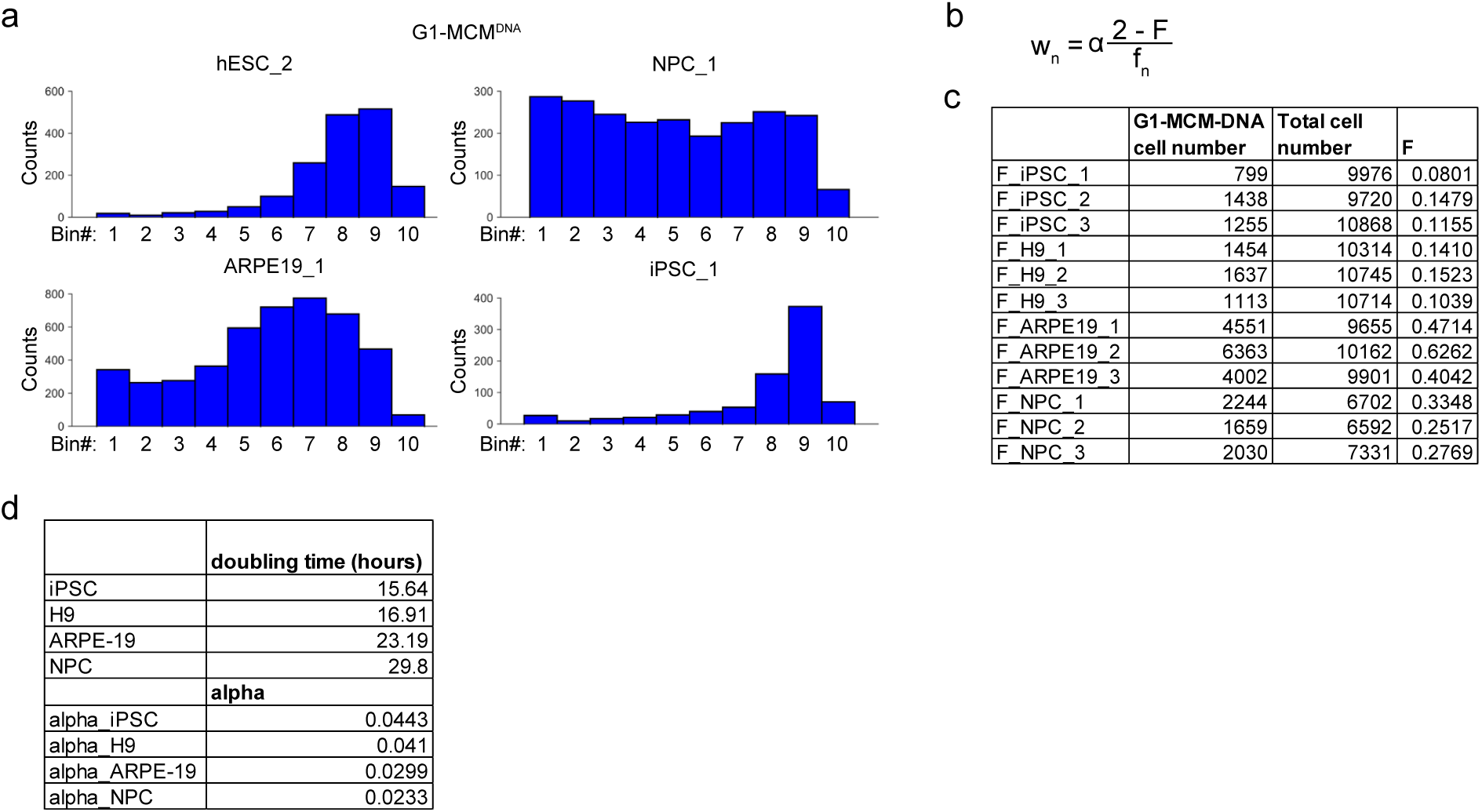
Ergodic Rate Analysis binning. (a) Histograms from G1-MCM^DNA+^ samples in Figure 2B, binned into 10 regions for ergodic rate analysis (see Methods). (b) Ergodic rate equation based on ^33^. w^n^ is the rate in each bin n, α is a constant accounting for doubling time, F is the fraction of cells in G1-MCM^DNA+^ out of all cells, f^n^ is the fraction of cells in each bin n out of G1-MCM^DNA+^ (see Methods). (c) Value of F for 3 biological replicates for each cell line, used to calculate Figure 2c. (d) Doubling time and alpha values used to calculate rate in Figure 2c.

**Supplementary Figure 4.**
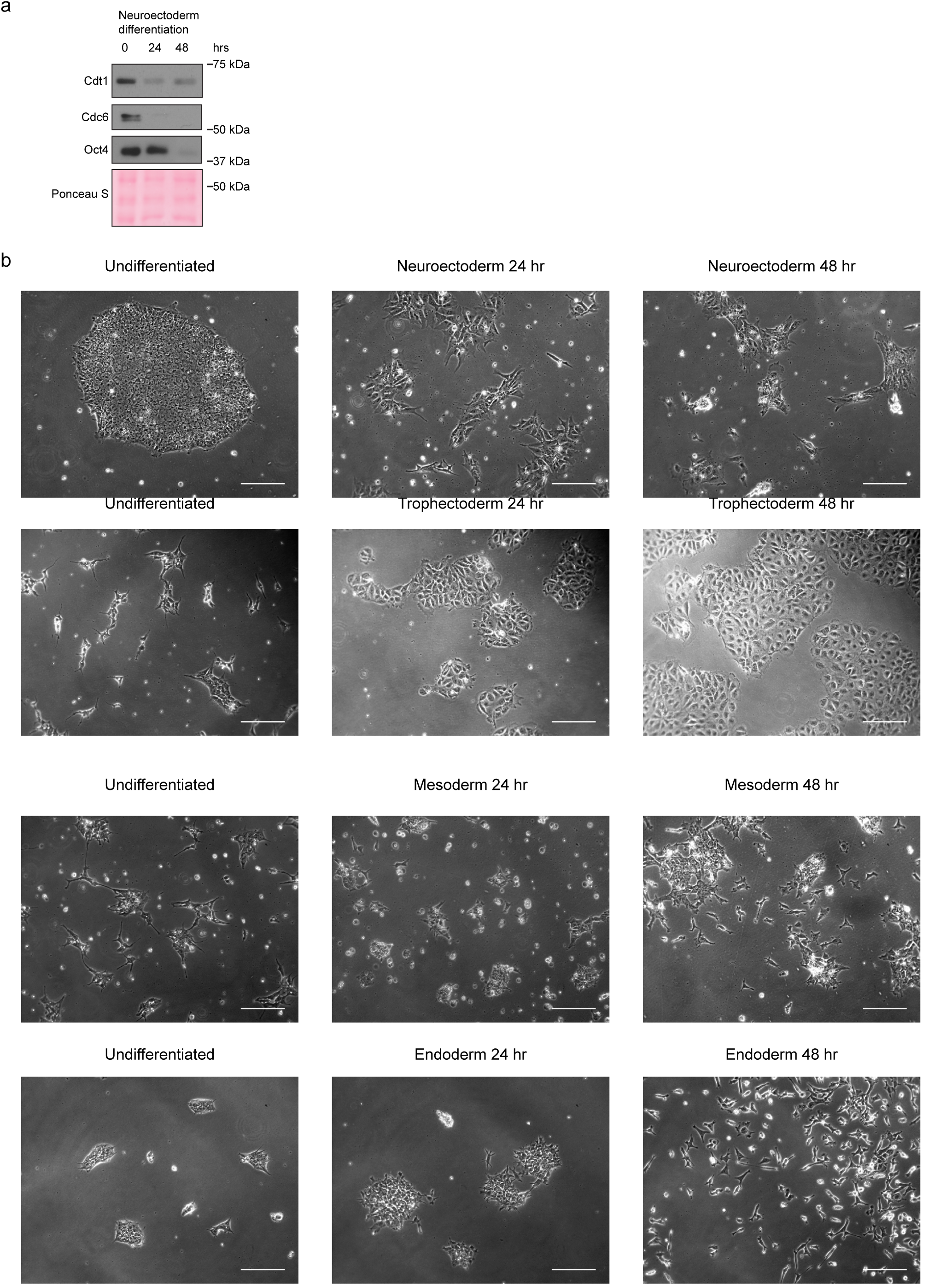
Stem cell differentiation. (a) Immunoblots of neuroectoderm differentiation, as in Figure 3a. (b) Representative phase contrast images in greyscale during the indicated differentiation protocols from Figure 3a. Scale bar is 50 μm.

**Supplementary Figure 5.**
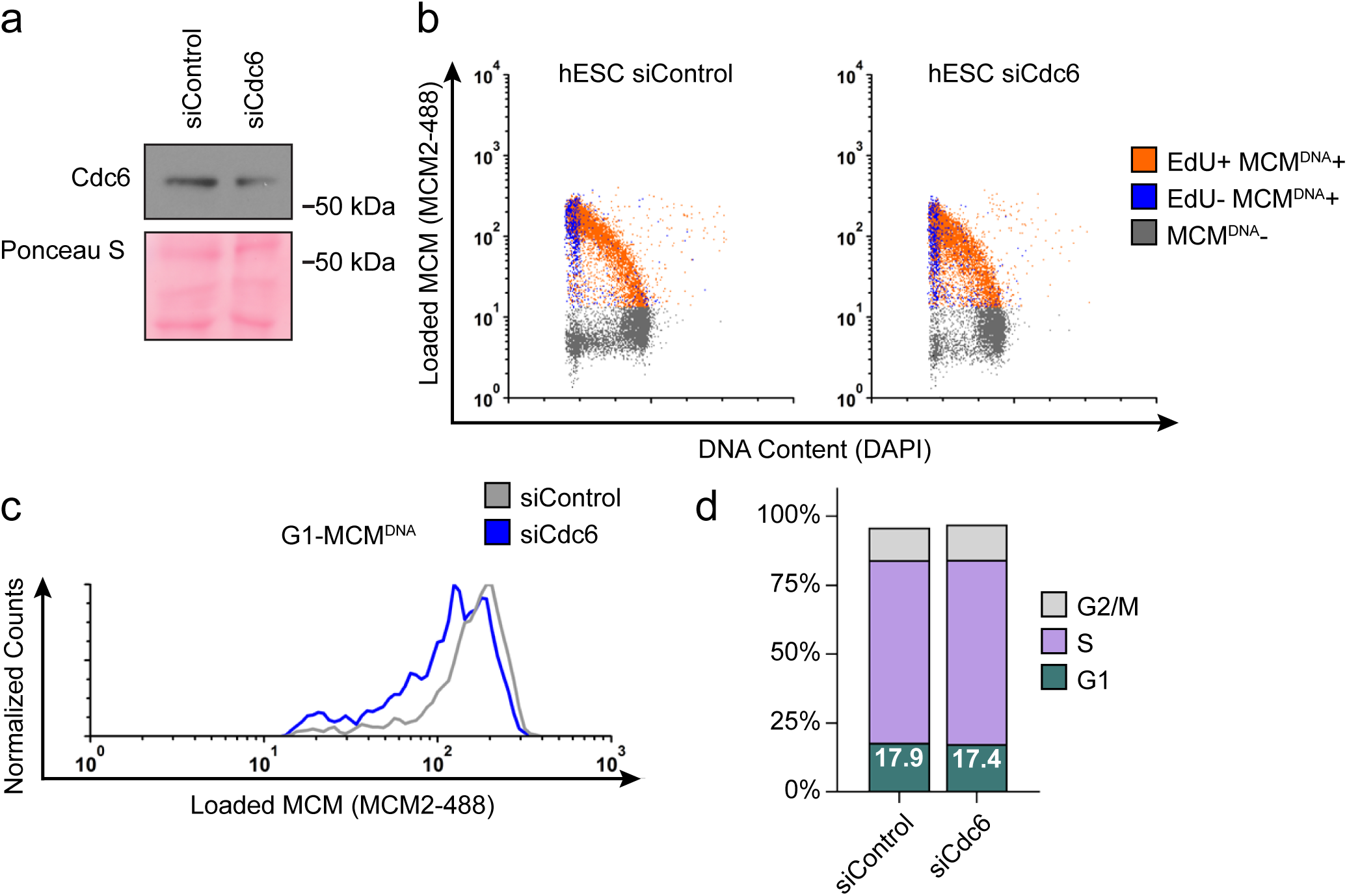
Reducing MCM loading rate by an alternative siRNA. (a) Immunoblot of total protein for hESCs treated with 100 nM siControl for 32 hr or 100 nM siCdc6 pool for 32 hr and pulse-labeled with EdU for 30 min prior to harvest. (b) Chromatin flow cytometry of cells from (a) stained with DAPI and anti-MCM2 and subjected to EdU detection. (c) Histogram MCM^DNA^ intensity in G1 cells. Counts for siCdc6 are normalized to the corresponding siControl sample. (d) Stacked bar graph of cell cycle distributions for samples in (a).

**Supplementary Figure 6.**
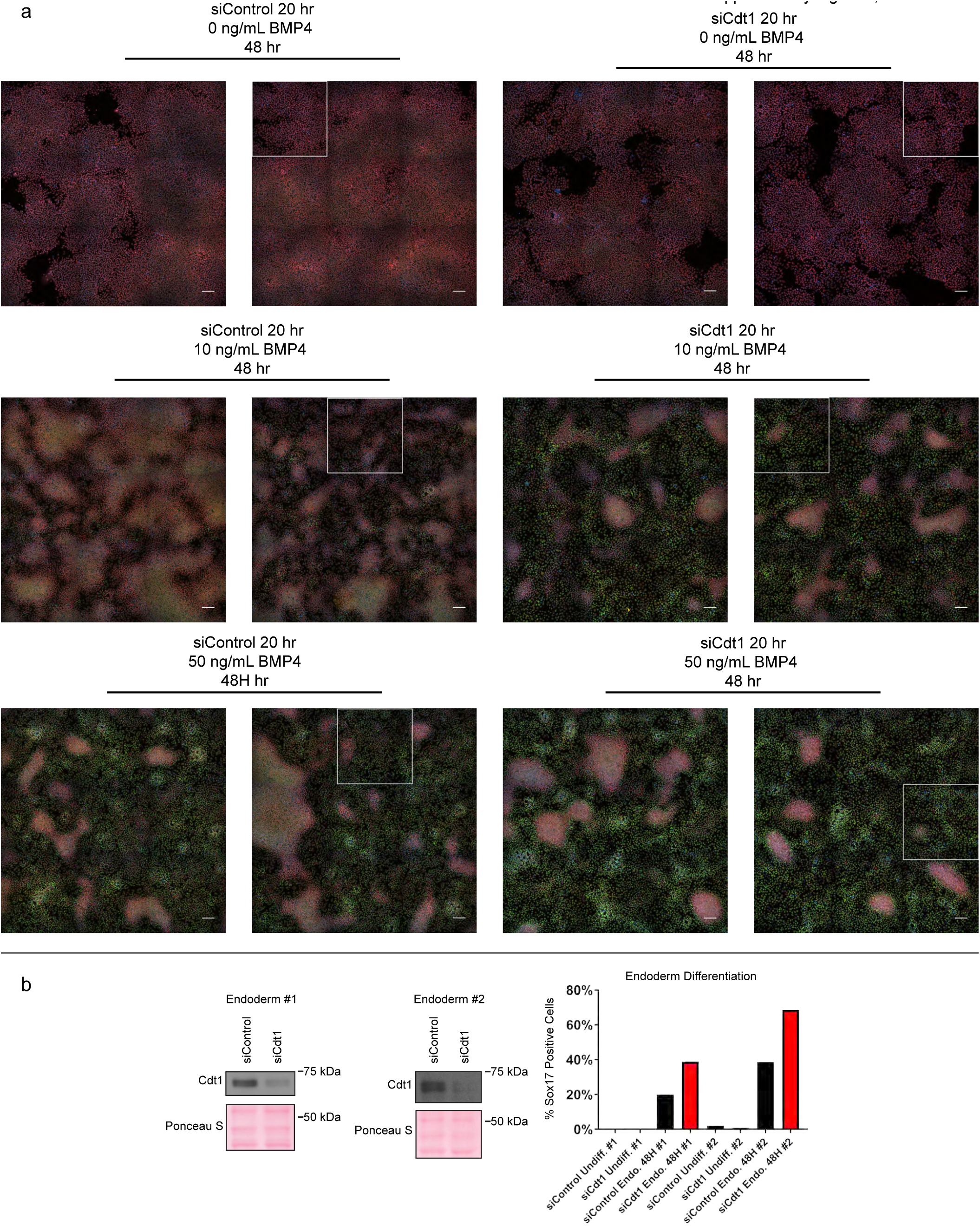
Complete microscopy dataset and endoderm differentiation. (a) Complete immunofluorescence microscopy data for Figure 7. White boxes mark the areas shown in Figure 7b; scale bars are 100 μM. (b) Immunoblots for Cdt1 in whole cell lysates of hESCs treated with 100 nM siControl or siCdt1 for 24 hr prior to initiating differentiation toward endoderm, two biological replicates. Bar graph indicates the percentage of Sox17-positive cells, n > 2,300 cells per condition.

**Supplementary Figure 7.**
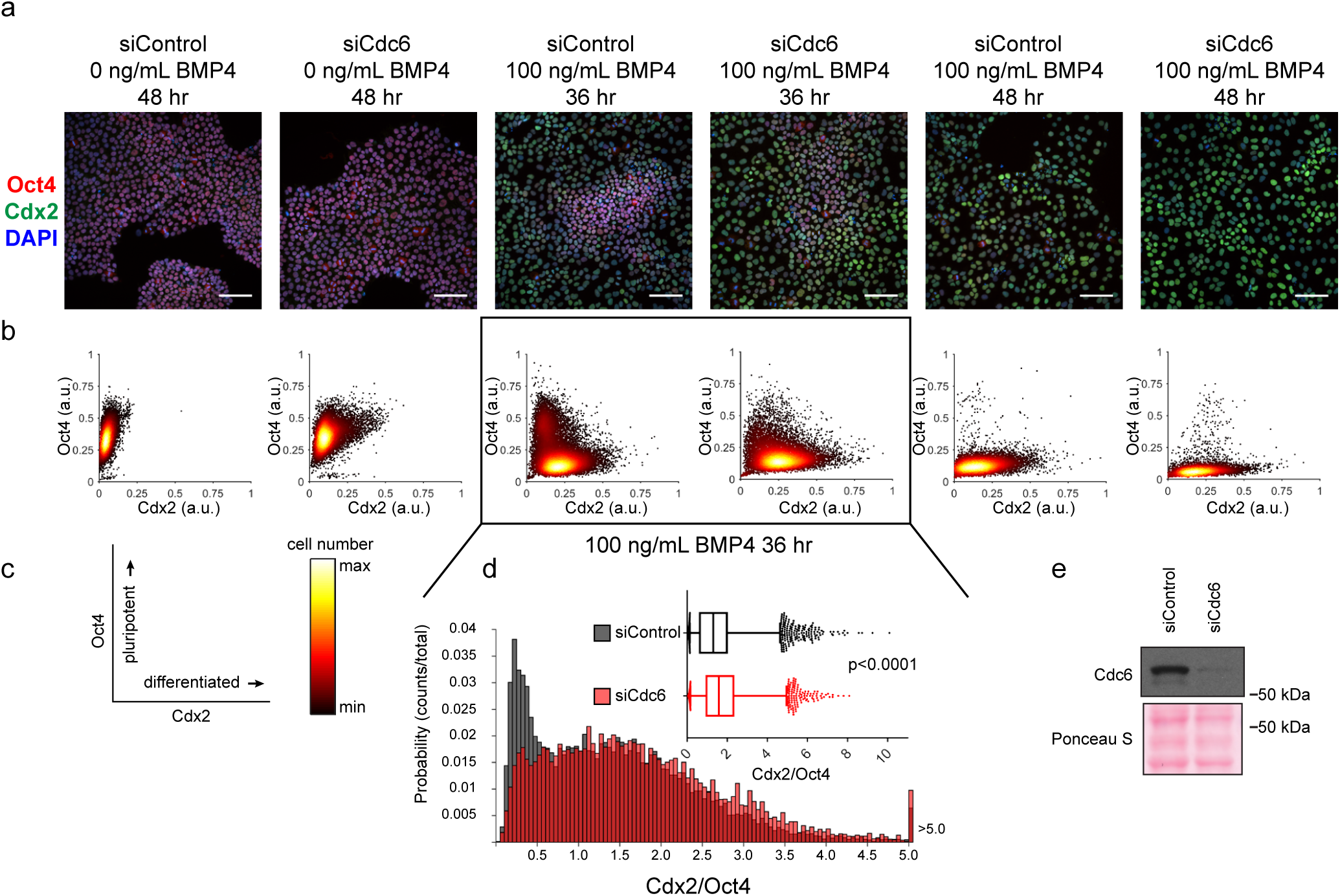
Slow MCM loading promotes differentiation. (a) Immunofluorescence microscopy of hESCs treated with 100 nM of siControl or 100 nM of siCdc6 for 32 hr and then treated with BMP4 as indicated. Cells were fixed and stained with DAPI (blue), Cdx2 antibody (green), and Oct4 antibody (red). Images are one region of 27 fields of view per condition; scale bar is 100 μm. (see methods). (b) Density scatterplots of mean fluorescence intensity (arbitrary units) of Oct4 and Cdx2 staining for each cell in each condition, >7,900 cells were quantified per condition. (c) Diagram of the relationship between Oct4 and Cdx2 in pluripotent and differentiated cells as plotted in (b); color bar for scatterplots in (b). (d) Histogram of the mean fluorescence intensity ratio Cdx2/Oct4 for all cells in siControl and siCdc6 treated with 100 ng/mL BMP4 for 36 hrs. Rightmost histogram bin contains all values greater than 5.0. the inset is a box-and-whiskers plot of the same data, center line is median, outer box edges are 25^th^ and 75^th^ percentile, whiskers edges are 1^st^ and 99^th^ percentile, individual data points are lowest and highest 1%, respectively. Medians are 1.296, 1.58, and means are 1.433, 1.728 for siControl and siCdc6, respectively. Samples compared by two tailed Mann-Whitney test,****p<0.0001. (e) Immunoblot for Cdc6 in whole cell lysates at 32 hr of siRNA treatment, prior to BMP4 treatment.

